# Single cell transcriptomics reveals a signaling roadmap coordinating endoderm and mesoderm diversification during foregut organogenesis

**DOI:** 10.1101/756825

**Authors:** Lu Han, Praneet Chaturvedi, Keishi Kishimoto, Hiroyuki Koike, Talia Nasr, Kentaro Iwasawa, Kirsten Giesbrecht, Phillip C Witcher, Alexandra Eicher, Lauren Haines, Yarim Lee, John M Shannon, Mitsuru Morimoto, James M Wells, Takanori Takebe, Aaron M Zorn

## Abstract

Visceral organs, such as the lungs, stomach, liver and pancreas, are derived from the fetal foregut through a series of inductive interactions between the definitive endoderm (DE) and the surrounding splanchnic mesoderm (SM). While patterning of DE lineages has been fairly well studied, paracrine signaling controlling SM regionalization and how this is coordinated with the epithelial identity during organogenesis is obscure. Here we used single cell transcriptomics to generate a high-resolution cell state map of the embryonic mouse foregut. This uncovered an unexpected diversity in the SM cells that developed in close register with the organ-specific epithelium. From these data, we inferred a spatiotemporal signaling roadmap of the combinatorial endoderm-mesoderm interactions that orchestrate foregut organogenesis. We validated key predictions with mouse genetics, showing the importance of endoderm-derived signals in mesoderm patterning. Finally, leveraging the signaling road map we generated different SM subtypes from human pluripotent stem cells (hPSCs), which previously have been elusive. The single cell data can be explored at: https://research.cchmc.org/ZornLab-singlecell.

## INTRODUCTION

In early fetal development, between embryonic day (E) 8.5 and E9.5 in mouse, equivalent to 17- 23 days of human gestation, a series of inductive tissue interactions between the definitive endoderm (DE) and the surrounding splanchnic mesoderm (SM) progressively patterns the naïve foregut tube into different progenitor domains. These domains further develop into distinct visceral organs including the trachea, lung, esophagus, stomach, liver, pancreas and proximal small intestine^1, 2^. The DE gives rise to the epithelial lining and parenchyma of the respiratory and digestive organs, while the SM gives rise to the mesenchymal tissues such as smooth muscle, fibroblasts and mesentery surrounding the visceral organs ^1,2^. This foregut patterning defines the landscape of the thoracic and abdominal cavities, setting the relative position of different organs. Disruptions in this process can lead to life threatening congenital birth defects.

A critical inductive role for the mesenchyme in gut tube organogenesis was first established in the 1960, when it was shown that SM transplanted from different anterior-posterior (A-P) regions of the embryo could instruct the adjacent epithelium to adopt the organ identity consistent with the original SM position^3^. Since that time we have learned much about the mesoderm derived paracrine signals in endoderm organogenesis ^1, 2^. However, most studies to date have focused on individual organ lineages or individual signaling pathways, and thus we still lack a comprehensive understanding of the temporally dynamic combinatorial signaling in the foregut microenvironment that orchestrates organogenesis. Moreover, several fundamental questions about the mesoderm have remained unanswered over the decades. How many types of SM are there, and does each fetal organ primordia have its own specific mesenchyme? How are the SM and DE lineages coordinated during organogenesis? What role if any does the endoderm have in regionalization of the mesoderm?

Initial specification and patterning of the embryonic mesoderm and endoderm occurs during gastrulation, from E6.25 to E8.0 in the mouse, as these germ layers progressively emerge from the primitive streak. The lateral plate mesoderm emerges from the streak after the extra-embryonic mesoderm, and is followed by the intermediate, paraxial and axial mesoderm^4, 5^. Concomitantly DE cells also delaminate from the streak and migrate along the outer surface of the mesoderm eventually intercalating into the overlaying visceral endoderm. By E8.0, morphogenetic processes begin to transform the bi-layered sheet of endoderm and mesoderm into a tube structure as the anterior DE folds over to form the foregut diverticulum and the adjacent lateral plate mesoderm containing cardiac progenitors migrates towards the ventral midline ^6^. The lateral plate mesoderm further splits into an outer somatic mesoderm layer next to the ectoderm which gives rise to the limbs and body wall, and an inner splanchnic mesoderm layer, which surrounds the epithelial gut tube ^7, 8^. The first molecular indication of regional identity in the SM is the differential expression of *Hox* genes along the A-P axis of the embryo^9^. However, in contrast to heart development, where cell diversification has been well studied^10-12^, the molecular mechanism governing the foregut SM regionalization are obscure, particularly during the critical 24 hours when the foregut DE subdivide into distinct organ primordia.

Recently, single cell transcriptomics have begun to examine organogenesis at an unprecedented resolution ^13-16^, however, studies in the developing gut have either primarily examined the epithelial component or later fetal organs after they have been specified ^17-19^. Here we used single cell transcriptomics of the mouse embryonic foregut to infer a comprehensive “cell state” ontogeny of DE and SM lineages, discovering an unexpected diversity in SM progenitor subtypes that develop in close register with the organ-specific epithelium. Projecting the transcriptional profile of paracrine signaling pathways onto these lineages, we inferred a roadmap of the reciprocal endoderm-mesoderm inductive interactions that coordinate organogenesis. We validated key predictions with mouse genetics showing that differential hedgehog signaling from the epithelium patterns the SM into gut tube mesenchyme versus mesenchyme of the liver. Leveraging the signaling road map, we generated different subtypes of human SM from hPSCs, which previously have been elusive.

## RESULTS

### Single cell transcriptomes define progenitor diversity in the developing foregut

To comprehensively define lineage diversification during foregut organogenesis, we performed single cell RNA sequencing (scRNA-seq) of the mouse embryonic foregut at three time points that span the period of early patterning and lineage induction: E8.5 (5-10 somites, ‘s’), E9.0 (12- 15s) and E9.5 (25-30s) (Fig. 1a, b). We micro-dissected the foregut between the posterior pharynx and the midgut, pooling tissue from 15-20 embryos for each time point. At E9.5, we isolated anterior and posterior regions separately, containing lung/esophagus and liver/pancreas primordia, respectively. A total of 31,268 single-cell transcriptomes passed quality control measures with an average read depth of 3,178 transcripts/cell. Cells were clustered based on the expression of highly variable genes across the population and visualized using uniform manifold approximation projection (UMAP) and t-distributed stochastic neighbor embedding (t-SNE) (Fig. 1c; Supplementary Fig. S1). This identified 24 cell clusters that could be grouped into 9 major cell lineages based on well-known marker genes: DE, SM, cardiac, other mesoderm (somatic and paraxial), endothelium, blood, ectoderm, neural crest and extraembryonic (Supplementary Fig. S1). DE clusters (4,448 cells) were characterized by co-expression of *Foxa1/2, Cdh1* and/or *Epcam*, whereas SM (10,097 cells) was defined by co-expression of *Foxf1* (Fig. 1d), *Vim* and/or *Pdgfra* as well as being negative for cardiac and other mesoderm specific transcripts.

**Fig. 1.**
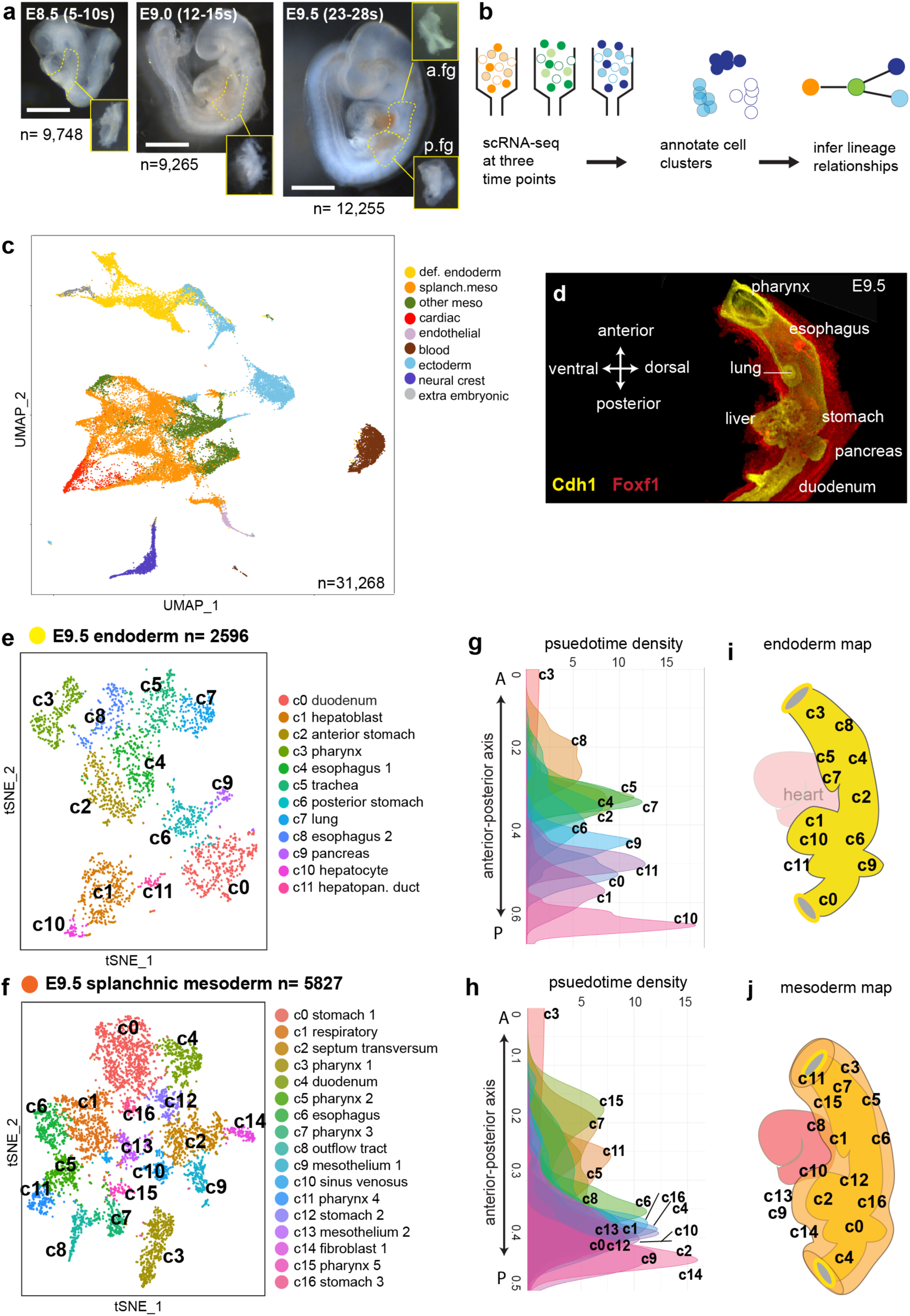
Single cell analysis of the mouse foregut endoderm and mesoderm lineages. **a**, Representative mouse embryo images at three developmental stages showing the foregut region (dashed) that was microdissected (insets) to generate single cells. At E9.5, anterior foregut (a.fg) and posterior foregut (p.fg) were isolated separately. E, embryonic day; s, somite number; n, number of cells. Scale bar 1 mm. **b**, Schematic of the RNA-seq workflow. **c**, UMAP visualization of 31,268 cells isolated from pooled samples of all three stages. Cells are colored based on major cell lineages. **d**. Whole-mount immunostaining of an E9.5 mouse foregut, showing the Cdh1+ endoderm and the surrounding Foxf1+ splanchnic mesoderm. **e** and **f**, t-SNE plot of *in-silico* isolated E9.5 endodermal (**e**) and splanchnic mesodermal (**f**) cells. **g** and **h**, Pseudo-spatial ordering of E9.5 endodermal (**g**) and mesodermal (**h**) cells along the anterior-posterior (A-P) axis. **i** and **j**, Schematic of the predicted locations of E9.5 cell types mapped onto **i**) the embryonic mouse foregut endoderm (yellow) and **j**) mesoderm (orange). def. definitive; meso, mesoderm; lg, lung; eso, esophagus; lv, liver; splanch; splanchnic. stm, septum transversum mesenchyme; sto, stomach; pha, pharynx. Source data for g,h are provides in the Source Data file.

To pinpoint lineage diversification in the DE and SM, we selected these cells *in silico* for further analysis. We defined 11 major DE clusters consisting of 26 stage-specific sub-clusters (E9.5, 12 clusters; E9.0, 8 clusters; E8.5, 6 clusters) and 13 major SM groups comprised of 36 stage-specific sub-clusters (E9.5, 17 clusters; E9.0, 12 clusters; E8.5, 7 clusters) (Fig. 1e, f, Supplementary Fig. S2 and S3, Table S1). We annotated clusters by comparing their distinguishing genes with published expression patterns of over 160 genes in the Mouse Genome Informatics (MGI) database ^20^. These data provide a comprehensive single cell resolution view of early foregut organogenesis and can be explored at URL: https://research.cchmc.org/ZornLab-singlecell.

Our annotations identified all the major DE organ lineages at E9.5 including: *Tbx1*+ pharynx, two *Nkx2-1/Foxa2*+ respiratory clusters, two *Sox2*+ esophagus clusters, two *Sox2/Osr1*+ stomach clusters, two *Alb/Prox1/Afp*+ hepatic clusters (c1_hepatoblasts and c10_ early hepatocytes with higher *Alb/HNF4a* expression), *Sox17/Pdx1*+ hepatopancreatic duct, *Pdx1/Mnx1*+ pancreas and *Cdx2*+ duodenum (Fig. 1e). Consistent with our dissections we did not detect any *Nkx2-1*+*/Hhex*+ thyroid progenitors. Similar to recent scRNA-seq analysis of the E8.75 gut epithelium ^19^, we also annotated half a dozen distinct DE progenitor states between E8.5 and E9.0, based on the restricted expression of lineage specifying transcription factors (TF), including *Otx2*+ anterior foregut, *Sox2/Sp5*-enriched dorsal lateral foregut, *Osr1/Irx1*-enriched foregut, *Hhex*+ hepatic endoderm, *Nkx2-3*+ ventral DE adjacent to heart and a small population of *Cdx2*+ midgut cells (Supplementary Fig. S2 and Table S1).

### Validation of novel mesenchymal subtypes

At all stages, the SM cell type diversity in the foregut was surprisingly complex, much more than previously appreciated (Fig. 1f and Supplementary Fig. S2). However, unlike the DE, SM populations were typically defined not just by one or two markers, but rather by a combination of multiple transcripts (Fig. 2a,b and Table S1). *In situ* hybridization and immunostaining of E9.5 foreguts and embryo sections confirmed that combinations of co-expressed transcripts defined different organ-specific SM subtypes (Fig. 2c-q and Supplementary Movies 1 and 2). The 17 SM cell populations at E9.5 included five *Tbx1/Prrx1*+ pharyngeal clusters, *Isl1/Mtus2*+ cardiac outflow tract cells, *Nkx6-1/Gata4/Wnt2*+ respiratory and *Nkx6-1/Sfrp2/Wnt4*+ esophageal mesenchyme (Fig. 2b-j). We annotated three *Barx1/Hlx*+ stomach mesenchyme populations, (one was probably ventral based on *Gata4* expression) and one *Hand1/Hoxc8*+ duodenum mesenchyme. We were unable to identify pancreas-specific mesenchyme and suspect that these cells were in the stomach or duodenum clusters (Fig. 2p,q).

**Fig. 2.**
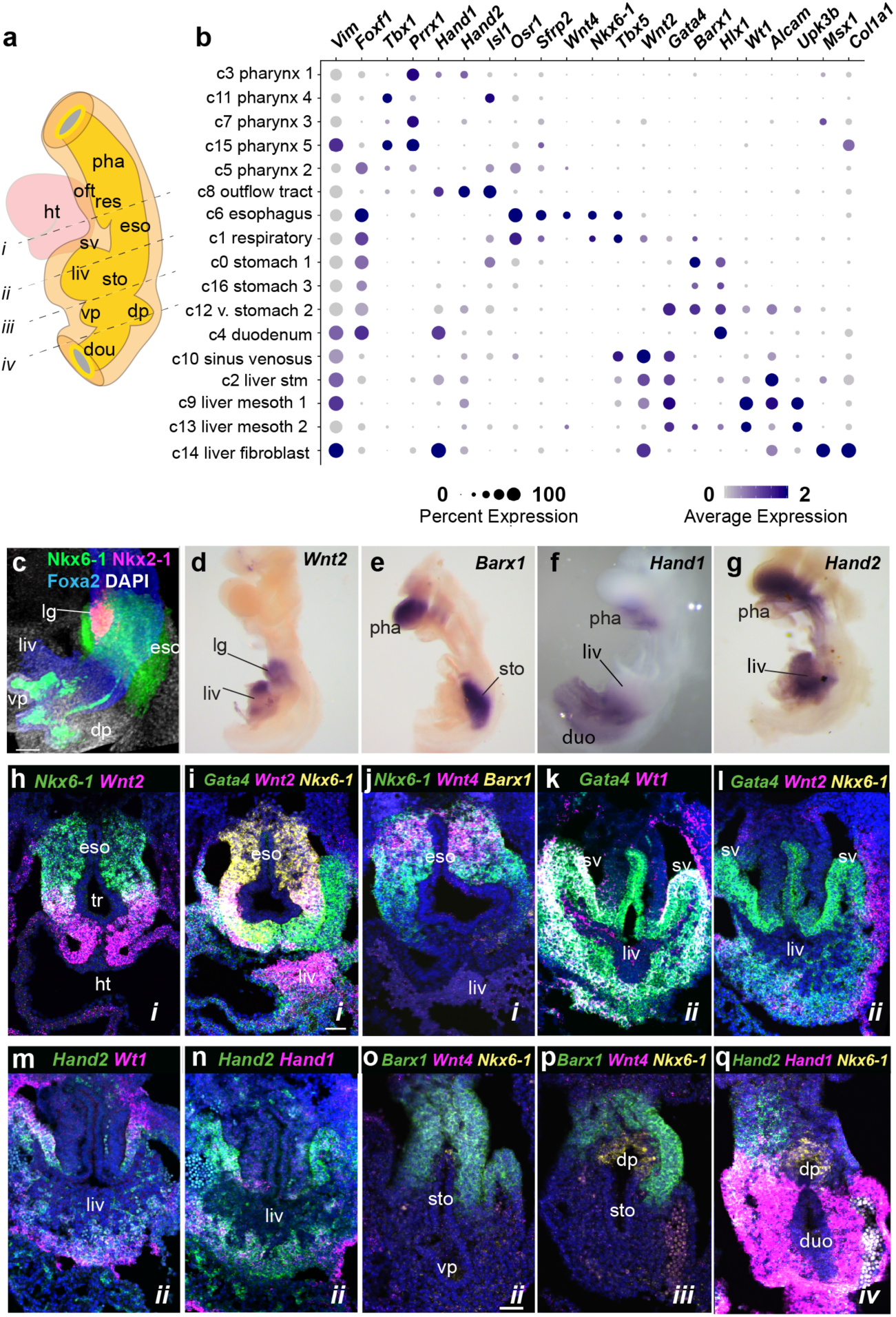
Lineage-restricted gene expression in different SM cell types. **a**, Schematic of the E9/5 foregut indicating the level of sections. **b**, Dot plot showing scRNA-seq expression of marker genes in different E9.5 SM cell clusters. **c-g**, Whole-mount immunostaining **c**) or *in situ* hybridization **d-g**) of dissected E9.5 foregut tissue. Scale bar 100μm. **h-q**, RNA-scope *in situ* detection on transvers E9.5 mouse embryos sections (*i-iv* indicates the A-P level of the section in **a**). Scale bar 50μm. duo; duodenum, dp; dorsal pancreas, eso; esophagus, ht; heart, lg; lung, liv; liver, oft, outflow tract, pha pharynx, res; respiratory, stm; septum transversum, mesenchyme, sto; stomach, sv; sinus venosus, vp; ventral pancreas.

Unexpectedly, the liver bud had five distinct mesenchymal populations. Data mining of MGI and *in situ* validation allowed us to annotate an *Alcam/Wnt2/Gata4*-enriched stm, a *Tbx5/Wnt2/Gata4/Vsnl1*+ sinus venosus, a *Msx1/Wnt2/Hand1/Col1a1*+ fibroblast population and two *Wt1/Gata4/Uroplakin*+ mesothelium populations (Fig. 2k-n and Supplementary Fig. S4). Interestingly we observed the restricted expression of *Hand1* and *Hand2* in the posterior versus anterior liver bud (Supplementary Fig. S4b) and the mutually exclusive expression of *Msx1* from *Wnt2* and *Wt1* (Supplementary Fig. S4e-f). This indicated extensive compartmentalization of the early liver bud mesenchyme warranting future investigation.

### Pseudotime spatial ordering of foregut cells

Different organs form at precise locations along the anterior-posterior (A-P) axis of the gut. To assess whether this was reflected in the single cell transcriptional profiles, we employed a pseudotime analysis, which several groups have recently used to examine positional information of cells in a continuous field of embryonic tissue ^16, 19, 21^. To this end we analyzed the DE and SM cells at each stage using diffusion maps, a dimensional reduction method for reconstructing developmental trajectories ^22, 23^. Anchoring the most anterior pharyngeal cluster as a root, we plotted the pseudotime density distribution for each cluster based on transition probabilities from root cells to all other cells in the graph (see Methods). Remarkably, this ordered both the DE and SM cell populations according to their appropriate A-P position in the embryo, indicating that the analysis represents an unbiased proxy of pseudo-space (Fig. 1g-j; Supplementary Fig.S2). The data also indicated that at this time in development, cells in the embryonic gut tube exhibit a continuum of transcriptional signatures of which spatially adjacent cell types having more similar expression profiles than distant cell types. Indeed the E9.5 clusters from the anterior dissections were located in the anterior half of the pseudo-space continuum, compared to posterior tissue, confirming the robustness of the computational ordering. Finally, we examined *Hox* genes which are known to be expressed in a co-linear fashion along the A-P axis and accordingly we observed a progressive increase of posterior *Hox* paralog expression in more posterior clusters, particularly within the SM (Supplementary Fig. S5a).

Combining the pseudo-space analysis, MGI curations and *in situ* validation, we were able to map each DE and SM population to their approximate locations in the gut tube (Fig. 1i, j; Supplementary Fig. S2). This revealed that the SM diversity mirrored DE lineages, indicating their closely coordinated development from the very beginning of organogenesis.

### Transcription factor code of foregut endoderm and mesenchyme

DE organ lineages have historically been defined by the overlapping expression domains of a few key transcription factors (TFs) ^1, 2, 24^. While some regionally expressed TFs have been reported in the SM, the single cell RNA-seq data allowed us to define a comprehensive combinatorial code of differentially expressed TFs that distinguish different SM and DE subtypes (Supplementary Fig. S5c and Table S1). This revealed new lineage-restricted markers such as homeodomain TF Nkx6-1. Well known for its expression in the pancreatic endoderm (Fig. 2p) ^25, 26^, Nkx6-1 was also specifically expressed in the respiratory and esophageal mesoderm at E9.5 (Fig. 2b,c,h-j, and Supplementary Movies 1 and 2). This TF code will facilitate lineage tracing experiments and prompt studies testing their role in mesenchymal differentiation.

### Synchronized endoderm and mesenchyme lineage trajectories

The transcriptional cell state complexity of the DE and SM doubled in just 24 hours between E8.5 and E9.5, reflecting progenitors forming more specialized cell types. To examine the temporal dynamics of lineage diversification, we visualized the single cell data using *SPRING* (Fig. 3a, b), an algorithm that represents k-nearest neighbors in a force directed graph, facilitating analysis of developmental trajectories^27^. Both the DE and SM trajectories progressed from a continuum of closely related cell states at E8.5 to transcriptionally distinct cell populations at E9.5 (Fig. 3a, b and Supplementary Fig. S6), consistent with the transition from multipotent progenitors to organ specific lineages. Importantly the cell clusters defined by tSNE were well-preserved in SPRING (Supplementary Fig. S6), supporting the robustness of the clustering. One striking observation evident in the structure of the *SPRING* plots was the apparent coordination of SM and DE lineage diversification over the 24 hours.

**Fig. 3.**
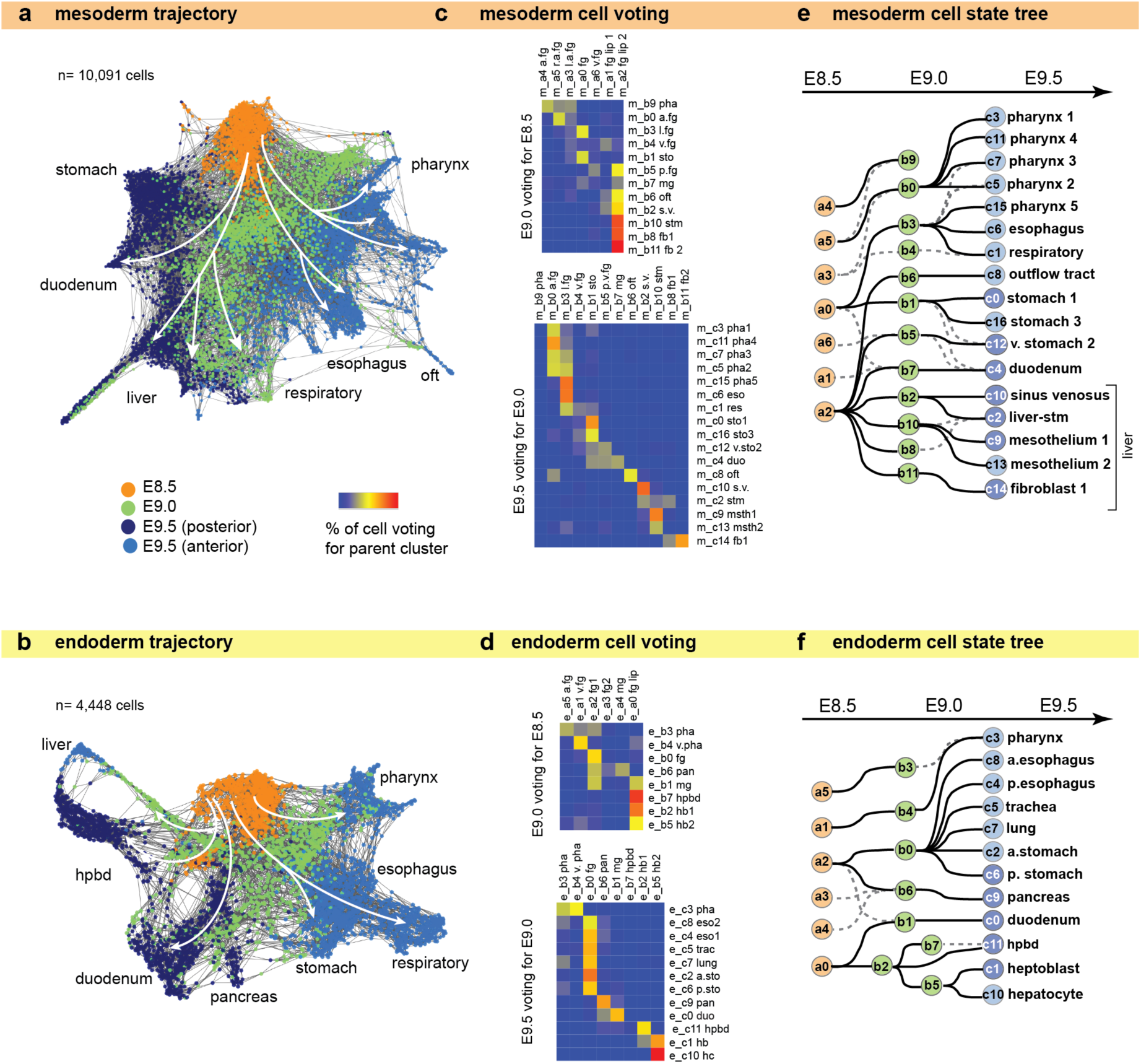
Coordinated endoderm and mesoderm cell trajectories. **a** and **b**, Force directed SPRING visualization of the **a**) splanchnic mesoderm (n=10,097) and **b**) definitive endoderm (**n**=4,448) cell trajectories. Cells are colored by developmental stage. White arrows indicate cell lineage progression. **c** and **d**, Confusion matrix summarizing “parent-child” single cell voting for **c**) SM and **d**) DE cells, used to construct the cell state tree. Each cell at the later time point (y-axis) voted for its most similar cell at the preceding time point (x-axis) based on transcriptome similarity (KNN) (see Methods). All of the votes for a give cluster are tabulated, normalized for cluster size (see Methods for details) and represented as a % of votes in the heatmap. E8.5, E9.0 and E9.5 clusters are designated as “a”, “b”, and “c”, respectively. **e** and **f**. Cell state trees of **e**) SM and **f**) DE lineages predicted by single cell voting. The top choice linking cell states of sequential time points are solid lines, and prominent second choices are dashed lines. Nodes are colored by stages and annotated with the cluster numbers.

To more clearly visualize the developmental trajectories associated with lineage diversification, we generated a consensus cell state tree using a single cell voting method, where each cell of a later time point votes for its most likely parent of the previous time point based on gene expression similarity. We then tabulate all the cell votes for each cluster (Fig. 3c,d) and represented this in a simple tree manifold (Fig. 3e,f). While we cannot rule out SM migration bringing distant cell types to a given organ, the data supported the notion of transcriptionally related cell states arising from the subdivision of common progenitor populations. Given that our time points were generated from pooled embryos of slightly different ages, it was possible that parent-child relationships could exist within a given time point. To address this and confirm the single cell voting results, we assessed each trajectory with a pseudotime analysis that computationally predicts progenitor states in a cell population (*Monocle*^*14*^). In general, the pseudotime analysis agreed with the single cell voting. But in the case of the liver endoderm, *Monocle* predicted a parent-child relationship within E9.0, where *Hhex*+ posterior foregut endoderm (cluster e_b2) gives rise to both *Prox1/Afp*+ hepatoblasts (e_b5) and *Prox1/Sox17/Pdx1*+ hepatopancreatic biliary progenitors (e_b7) (Supplementary Fig. S7), consistent with *in vivo* lineage tracing experiments ^28, 29^.

Overall the DE trajectories inferred by the single cell transcriptomes are consistent with experimentally determined fate maps ^28-30^, demonstrating the robustness of our analysis and suggesting that the SM trajectories, which previously have not been well defined, may also represent lineage relationships. Having said that we caution that cells with this similar transcriptomes may not necessarily be lineage-related. Indeed there are cases where cells from different lineages such as ventral and dorsal pancreas can converge on similar transcriptional profiles. Thus our results establish a theoretical framework for future experimental analysis of foregut mesenchyme development.

### Coordinated development of multi-potent progenitors

A close examination of the DE and SM trajectories suggests the coordinated development from multipotent progenitors within adjacent endoderm and mesoderm tissue layers. For example, at E8.5 the DE lateral foregut cells (e_a2) and the spatially neighboring SM cells (m_a0) both express the TF *Osr1*, and the trajectories predict that these two cell populations are multipotent progenitors, giving rise to the respiratory, esophageal and gastric epithelium and mesenchyme respectively (Fig. 4a-b). As development proceeds, different cell populations appear to be segregated as they progressively express distinct lineage regulating TFs and growth factors (Fig. 4a-d). *In situ* validation confirmed that *Osr1* is expressed in both the epithelium and mesenchyme of the presumptive esophagus, lung and stomach at E9.5 (Fig. 4e-g).

**Fig. 4.**
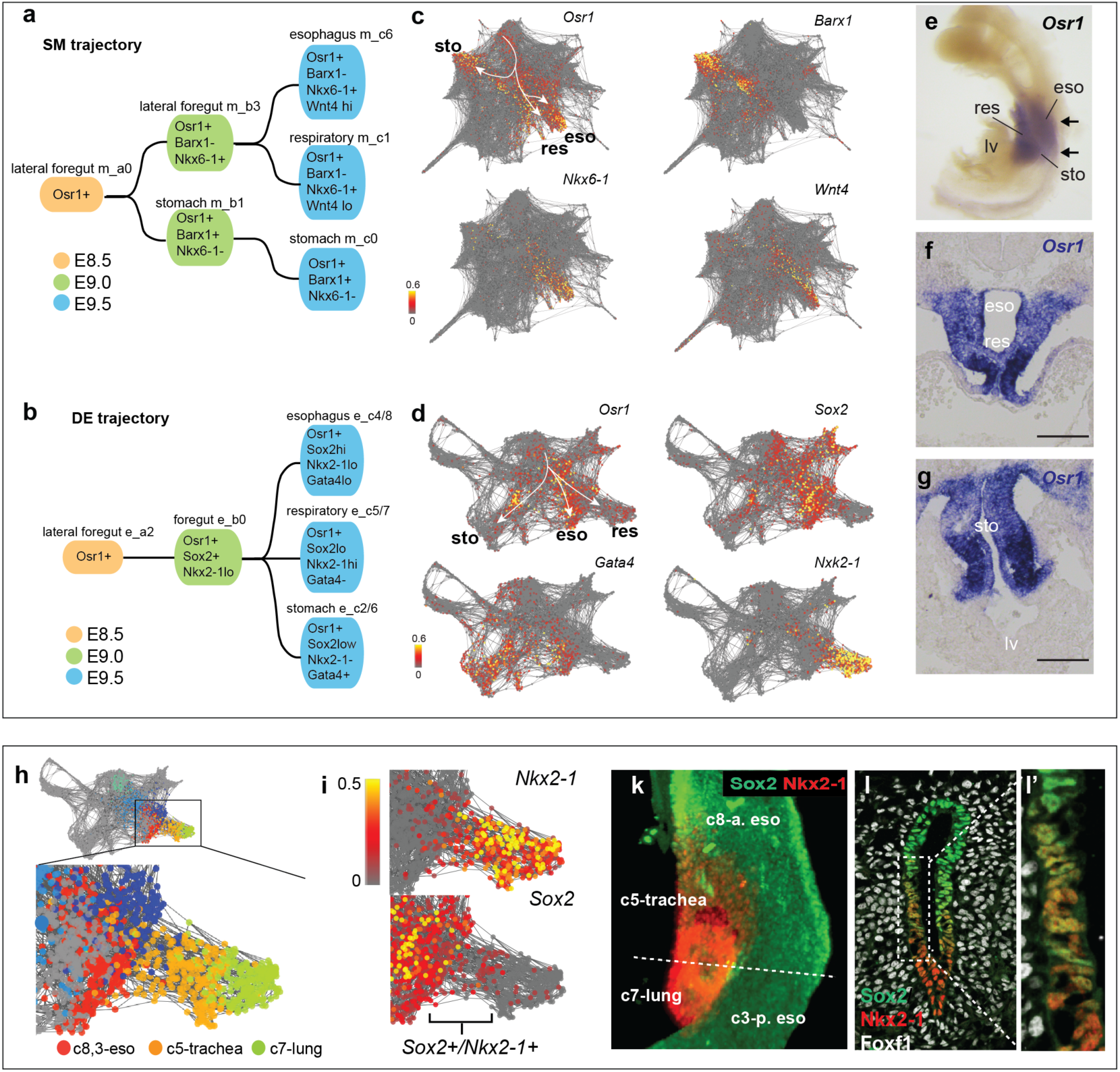
Coordinated development of multipotent progenitors. **a** and **b**, Graphical illustration of the esophageal-respiratory-gastric cell state trajectories for **a**) SM and **b**) DE with key marker genes. This suggests the coordinated development of *Osr1*+ multi-lineage progenitors. **c** and **d**, SRPING plots of c) SM and d) DE projecting the expression of key genes. **e**, *in situ* hybridization of *Osr1* in dissected foregut, showing *Osr1* is expressed in the respiratory, esophageal and gastric regions. **f** and **g**, *in situ* hybridization of *Osr1* in sections across the respiratory and gastric regions within the foregut, showing that *Osr1* is expressed in both the endodermal and mesenchymal cells. **h**, SPRING plot of the DE esophageal-respiratory lineages **i**. *Nkx2-1* and *Sox2* expression are projected onto the SPRING plot, showing co-expression at the esophageal-tracheal boundary. **k**, Sox2 and Nkx2-1 whole mount immunostaining of a E9.5 mouse foregut. **l**, Sox2, Nkx2-1 and Foxf1 immunostaining of a transverse E9.5 foregut section, confirming a rare population of Sox2/Nkx2-1 co-expressing cells. **l’**, Higher magnification of box in l.

Furthermore, a close examination of the DE trachea cluster suggested a transitional cell population co-expressing the respiratory maker *Nkx2-1* and the esophageal marker *Sox2* at E9.5 when the foregut is being patterned along the dorsal-ventral axis (Fig. 4h-i). Immunostaining confirmed that this was indeed a rare Nkx2-1/Sox2+ cell population at the prospective tracheal-esophageal boundary (Fig. 4k-l), which recent studies have demonstrated to be critical in tracheoesophageal morphogenesis^31, 32^. In sum, the foregut lineage trajectories predicted from the single cell transcriptomes represent a valuable resource for further studies.

### Predicting a signaling road map of organ induction

We next sought to computationally predict the paracrine signaling microenvironment in the foregut that controls these cell fate decisions (Fig. 5a,b). We calculated metagene expression profiles for all the ligands, receptors and context-independent response genes in each DE and SM cluster for six major signaling pathways implicated in organogenesis: BMP, FGF, Hedgehog (HH), Notch, retinoic acid (RA) and canonical Wnt (see Methods & Supplementary Fig. S8, Table S2). Leveraging our spatial map of each cell population in the foregut (Fig. 1i,j) we ordered cell populations along the A-P axis such that DE and SM cell types most likely to be in direct contact were opposite one another in the signaling diagram (Fig. 5c). We then used the metagene expression levels to predict potential ligand-receptor pairs and the likelihood that a given cell population was responding to local paracrine or autocrine signals (Fig. 5a-c, Supplementary Fig. S9). We benchmarked the metagene expression thresholds on experimentally validated interactions from the literature. Also we limited potential ligand-receptor pairings to nearby cell clusters, consistent with the generally accepted view that these pathways act over a relatively short range. Together this analysis revealed a hypothetical combinatorial signaling network (Fig. 5a-c, Supplementary Fig. S9).

**Fig. 5.**
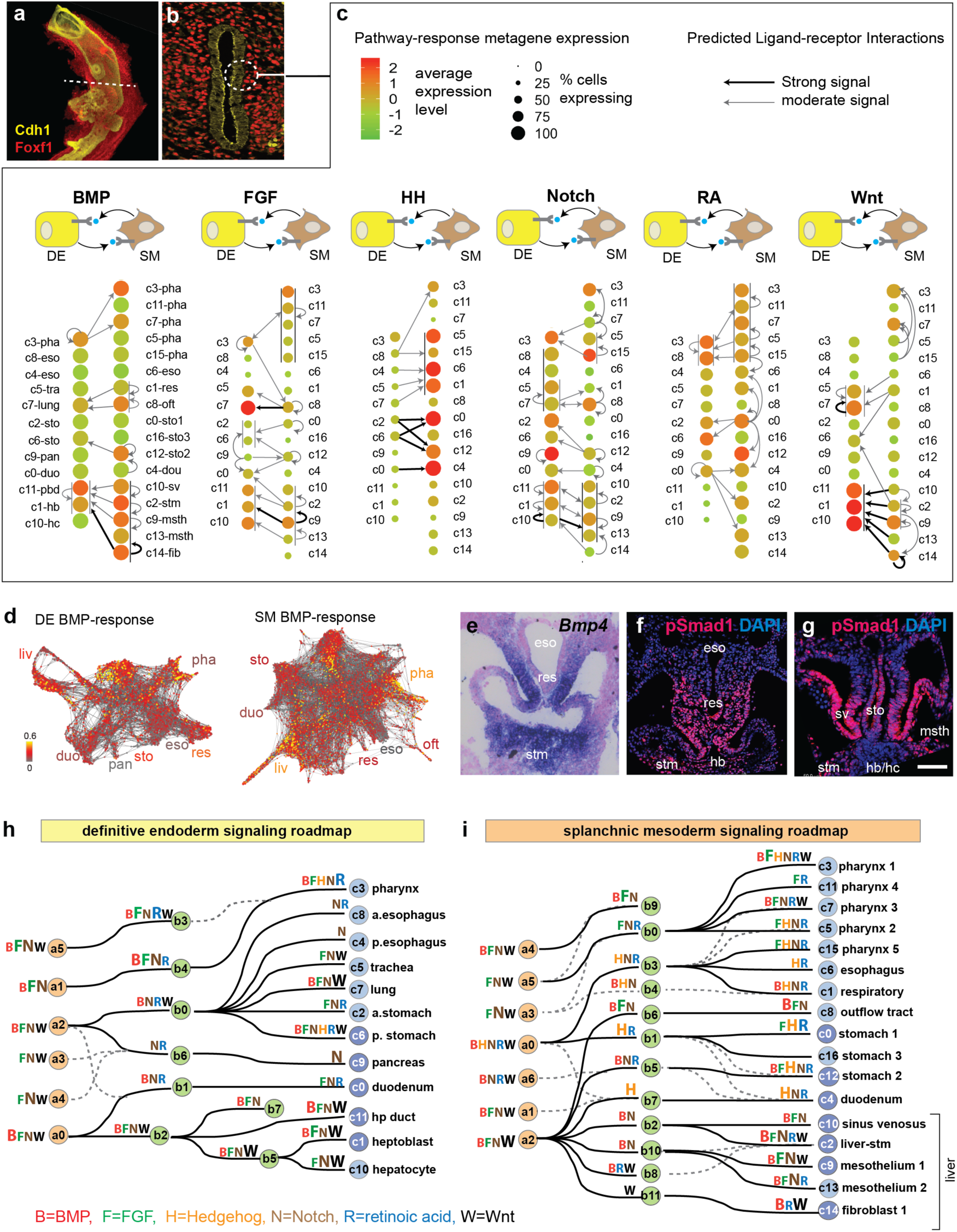
Computationally inferred receptor-ligand interactions predict a signaling roadmap of foregut organogenesis. **a** and **b**, E9.5 foregut immunostaining of Cdh1 (epithelium) and Foxf1 (mesenchyme) in **a**) whole mount (same image as Fig. 1d) and **b**) section, showing the epithelial-mesenchymal tissue microenvironment (dashed circle). **c**, Predicted receptor-ligand interactions between adjacent foregut cell populations. The schematics show paracrine signaling between the DE (yellow cells) and the SM (brown cells) for six major pathways. E9.5 DE and SM cell clusters are ordered along the anterior to posterior axis based to their locations *in vivo*, with spatially adjacent DE and SM cell types are across from one another. Colored circles indicate the relative pathway response-metagene expression levels, predicting the likelihood that a given cell population is responding to the growth factor signal. Thin vertical lines next to clusters indicate different cell populations in spatial proximity that are all responding to a particular signal pathway. Arrows represent the predicted paracrine and autocrine receptor-ligand interations (see Methods). **d**, BMP response-metagene expression levels projected on the DE and SM SPRING plot. **e**, *in situ* hybridization of *Bmp4* in a foregut transverse section, showing the expression of in the respiratory mesenchyme and the stm. **f** and **g**, pSmad1 immunostaining in foregut transverse sections, indicating BMP signal response in the respiratory and liver DE and SM. **h and i**, Signaling roadmap summarizing the inferred signaling state of all 6 pathways projected on the **h**) DE and **i**) SM cell state trees suggests the combinatorial signals predicted to control lineage diversification. The letters indicated the putative signals at each step, with larger font indicating a stronger signaling response. a, anterior; p, posterior; hp, hepatopancreatic; stm, septum transversum mesenchyme.

Overall the computational predictions are consistent with known expression patterns of ligands and receptors, and identified most known signaling interactions controlling DE lineage specification. This includes mesoderm derived BMP, FGF and Wnt promoting DE liver and lung fate, and autocrine notch signaling in the DE endocrine pancreas ^1, 2, 33, 34^. This suggested that previously undefined SM signaling predictions are also likely to be accurate. To test this we examined BMP signaling as an example. Consistent with the scRNA-seq data, *in situ* hybridization confirmed high levels of *Bmp4* ligand expression the stm and the respiratory mesenchyme, while immunostaining for phospho-Smad1/5/8, the cellular effector of BMP signaling, confirmed autocrine and paracrine signaling in the developing liver and respiratory mesenchyme and epithelium respectively as predicted (Fig. 5e-g).

We projected the signaling response-metagene expression levels onto the *SPRING* plots and cell state tree which revealed spatiotemporally dynamic signaling domains that correlated with cell lineages (Fig. 5d, Supplementary Fig. S10). In general, the transcriptome data predicts locally restricted interactions, with the SM being the primary source of BMP, FGF, RA and Wnt ligands, signaling to both the adjacent DE and within the SM itself (Fig. 5c). In contrast, HH ligands are produced by the DE and signal to the gut tube SM, with no evidence of autocrine activity in the DE (Fig. 5c). Combining the data for all six signaling pathways onto the cell state trees, we generated a comprehensive roadmap of the combinatorial signals predicted to coordinate the temporal and spatial development of each DE and SM lineage (Fig. 5h,i). This analysis predicts a number of previously unappreciated signaling interactions and represent a hypothesis generating resource for further experimental validation.

### Testing the role of epithelial Hedgehog signaling in foregut mesenchyme patterning

To genetically test the predictive value of the signaling roadmap, we focused on HH activity, which is suggested by the scRNA-seq to be high in gut tube SM (esophagus, respiratory, stomach and duodenum) but low in the pharyngeal and liver SM (Fig. 6a-c). HH ligands stimulate the activation of Gli2 and Gli3 TFs, which in turn promote the transcription of HH-target genes (e.g. *Gli1)*^35^. As expected, mouse embryo sections confirmed that *Shh* ligand was expressed in the gut tube DE with high levels of *Gli1-LacZ* expression in the adjacent SM. By contrast, the hepatic endoderm did not express *Shh* ^36^ and the hepatic SM had very few if any *Gli1-LacZ* positive cells (Fig. 6d). To define the function of HH in SM patterning, we performed bulk RNA-seq on foreguts from *Gli2- /-;Gli3-/-* double mutant embryos, which lack all HH activity and fail to specify respiratory fate ^34^. Comparing homozygous mutants to heterozygous littermates, we identified 156 HH/Gli-regulated transcripts (Fig. 6e; Supplementary Table. S3). Given the caveat that this bulk RNA sequencing is performed with both endoderm and mesoderm, we examined the enrichment of these HH-regulated transcripts in the transcriptome of DE and SM single cell clusters. This revealed that most transcripts were expressed the SM compared to the DE. Importantly, transcripts downregulated in Gli2/3-mutants (n=80) were normally enriched in the gut tube SM, whereas upregulated transcripts (n=76) were normally enriched in the liver or pharyngeal SM (Fig. 6e-g). Interestingly HH/Gli-regulated transcripts, including downregulated TFs (Osr1, Tbx4/5, Foxf1/2) and upregulated TFs (Tbx18, Lhx2 and Wt1), have been implicated in respiratory and hepatic development respectively (Fig. 6e; Supplementary Table S3) ^37^. This genetic analysis confirmed the predictive value of the signaling roadmap where differential HH activity promotes gut tube versus liver and pharyngeal SM (Fig. 5i), in part by regulating other lineage specifying TFs and signaling proteins.

**Fig 6.**
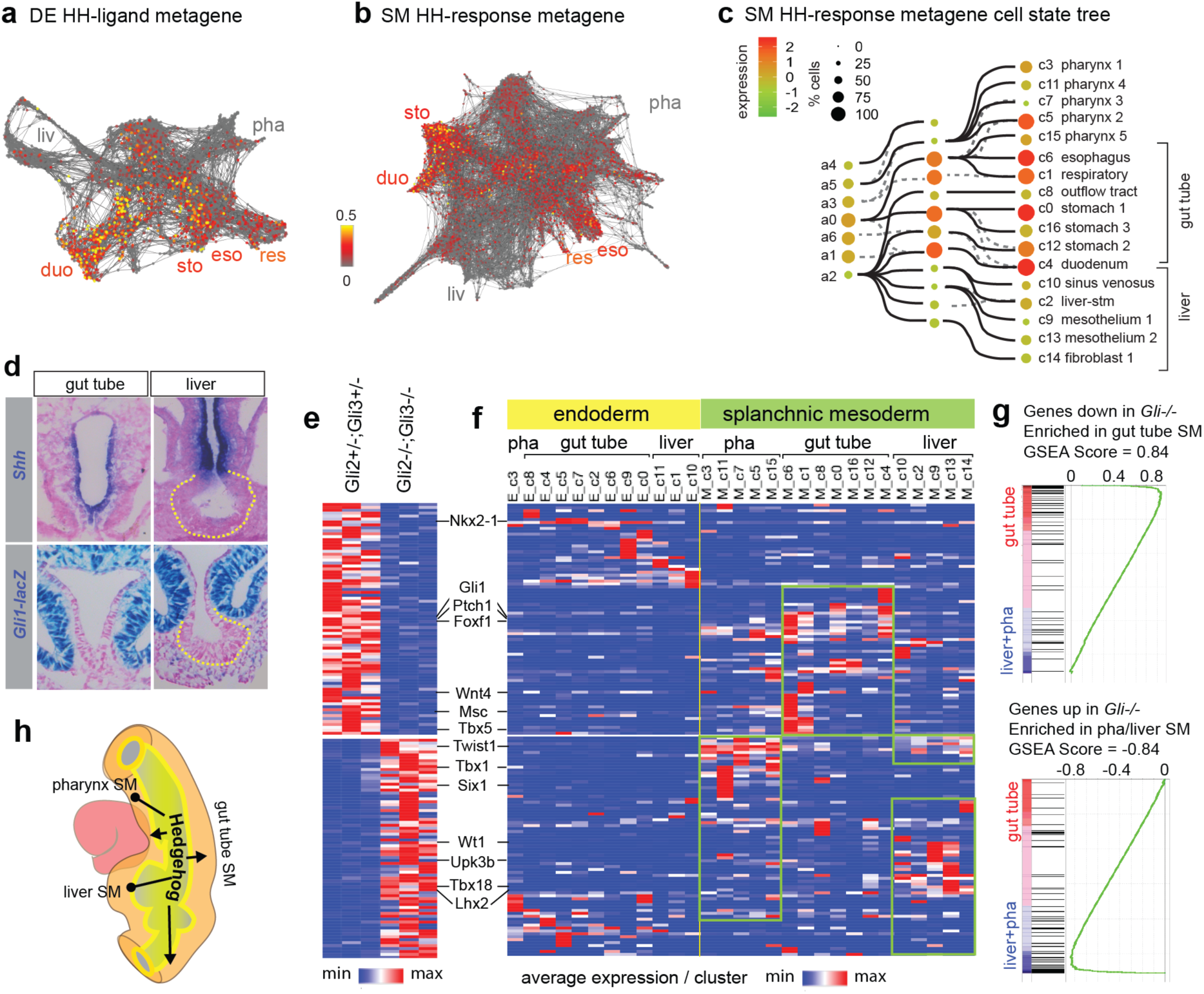
Genetic test of the signaling roadmap revealed that HH promotes gut tube versus liver mesenchyme. **a** and **b**, SPRING visualization of **a**) the HH ligand-metagene expression in DE cells and **b**) HH response-metagene expression in SM cells. **c**, The HH response-metagene expression projected onto the SM cell state tree, showing low HH activity in the liver and pharynx SM but high activity in the gut tube mesenchyme. **d**, *Shh* is expressed in the gut tube epithelium but not in the hepatic epithelium (outlined). *Gli1-lacZ*, a HH-response transgene, is active in the gut tube mesenchyme but not in the liver stm. **e**. Differentially expressed genes between *Gli2-/-, Gli3-/-* and *Gli2*+*/-, Gi3*+*/-* mouse E9.5 foreguts through bulk RNA-sequencing (log2 FC >1, FDR < 5%). **f**, Heatmap showing average expression of HH/Gli-regulated genes (from Fig 6**e**) in E9.5 DE and SM single cell clusters. **g**, Gene set enrichment analysis (GSEA) reveals specific cell type enrichment of HH/Gli-regulated genes. **h**, Schematic of HH activity in the foregut.

Our data, together with previous work, suggested a model where the reciprocal epithelial-mesenchymal signaling network coordinates DE and SM lineages during organogenesis. In this model, SM-derived RA induces a regionally restricted expression of Shh in the DE by E9.0 ^34^, which then signals back to the SM, establishing broad pharynx, gut tube and liver domains. Other SM ligands (BMP, FGF, Notch, RA and Wnt), with distinct combinations of regional expressions in these three broad domains, then progressively subdivide DE and SM progenitors in a coordinated manner. In the future it will be important to test this model by cell-specific genetic manipulations.

### Differentiation of splanchnic mesenchyme-like lineages from human PSCs

We next tested whether our new SM markers and signaling roadmap could be used to direct the differentiation of distinct SM subtypes from human pluripotent stem cells (hPSC), which to date have been elusive. Previous studies have established protocols to differentiate hPSC into lateral plate mesoderm (lpm) and cardiac tissue ^38^. Although both the SM and heart are derived from the lpm, the single cell data suggested that in the mouse, the early SM experiences more RA signaling than the early cardiac mesoderm. This was confirmed by RA-responsive *RARE:lacZ* transgene expression in E8.5 embryos (Supplementary Fig. S11a, b). Accordingly, addition of RA to the lpm differentiation media on days (d) 2-4 down-regulated the cardiac markers *NKX2-5, ISL1* and *TBX20* and promoted the SM markers *FOXF1, HOXA1, HOXA5 and WNT2* (Fig. 7b, Supplemental Fig. S11c,d). This is consistent with the mouse scRNA-seq data which shows that E8.5 SM expresses *Nkx2-5, Isl1* and *Tbx20*, at lower levels than the cardiac mesoderm. Examination of *PAX3, PRRX1* and *CD31* confirmed that the d4 SM cultures did not express significant levels of endothelial, somatic or limb mesenchyme markers (Supplemental Fig. S11c).

**Fig. 7.**
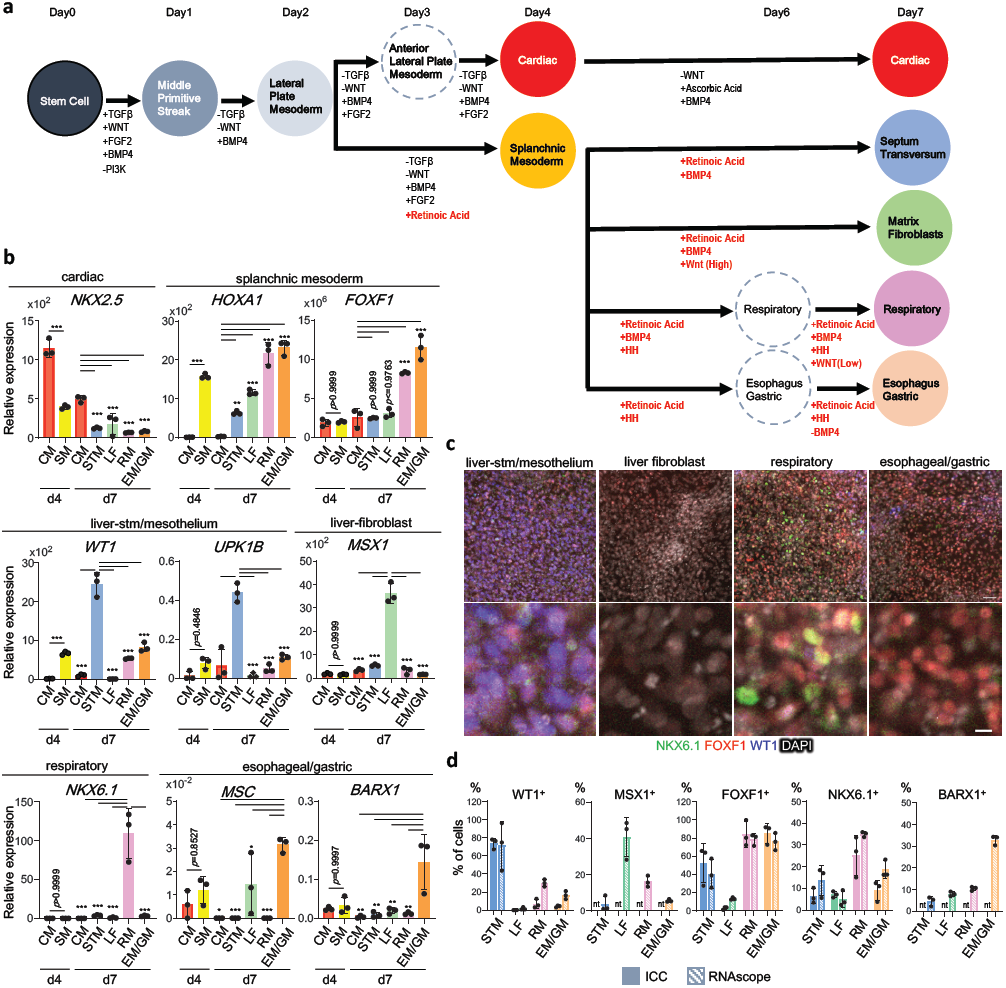
Generation of splanchnic mesoderm-like progenitors from human PSCs. **a**, Schematic of the protocol to differentiate hPSCs into SM subtypes. Factors in red indicate signals predicted from the mouse single cell signaling roadmap. **b**, RT-PCR of markers with enriched expression in specific SM subtypes based on the mouse single cell data: cardiac (*NKX2-5*), early SM (*FOXF1, HOXA1*); liver-stm/mesothelium (*WT1, UKP1B*), liver-fibroblast (*MSX1*), respiratory SM (*NKX6-1*+, *MSC*-), esophageal/gastric (*MSC, BARX1*). Columns show the means ± S.D. Tukey’s test *p<0.05, **p<0.005, ***p<0.0005. **c**, Immunostaining of Day 7 cell cultures. Scale bar; 50μm (*upper panels*), 10μm (*lower panels*). **d**, Quantification of % cells positive for the indicated immunostaining or RNA-scope in situ hybridization. Columns show the means ± S.D. (n=3). Tukey’s test, *p<0.05, **p<0.005, ***p<0.0005. Source data are provided as a Source Data file.

We next treated the primitive SM with different combinations of HH, RA, Wnt and BMP agonists or antagonist from d4-d7 (Fig. 7a), to drive organ-specific SM-like lineages based on the roadmap. As predicted, the HH-agonist promoted gut tube identity and efficiently blocked the hepatic fate. In the HH treated cultures, addition of RA and BMP4 (RA/BMP4) followed by WNT on d6-7 promoted gene expression consistent with respiratory mesenchyme (*NKX6-1, TBX5* and *WNT2*) with low levels of esophageal, gastric or hepatic markers. In contrast, addition of RA and BMP4-antagoist on d6-7 promoted an esophageal/gastric-like identity (*MSC, BARX1, WNT4* and *NKX3-2*) (Fig. 7b-c and Supplementary Fig. S12d). In the absence of HH agonist, cells treated with RA/BMP had a gene expression profile similar to liver stm and mesothelium (*WT1, TBX18, LHX2* and *UPK1B*), whereas RA/BMP4/WNT treated cells expressed liver-fibroblast markers (*MSX1/2* and *HAND1*). Immunostaining and RNA-scope confirmed the RT-PCR analysis (Fig. 7c-d and Supplementary Fig. S12a-c) showing that ∼70-80% cells in the liver stm/mesothelium-like cultures were WT1+,MSX1-,NKX6-1-, whereas the other populations appear to be around 30- 40%. The remainder of cells appeared to be undifferentiated rather than an alternative lineage. While future work is needed to optimize the culture conditions for each lineage, these data provided a proof of principle that the signaling roadmap inferred from the mouse scRNA-seq data can be used to direct the differentiation of different organ specific SM subtypes from hPSC.

## DISCUSSION

We have used single cell transcriptomics to define the complexity of DE and SM cell types in the embryonic mouse foregut over the first 24 hours of organogenesis as the primitive gut tube is subdivided into distinct organ domains. Our analysis revealed an unexpected diversity of distinct cell types in the foregut mesenchyme, defined by new marker genes and a combinatorial code of transcription factors. Cell trajectories indicate that the development of organ-specific DE and SM is closely coordinated, suggesting a tightly regulated signaling network. We computationally predicted a putative ligand-receptor signaling roadmap of the reciprocal epithelial-mesenchymal interactions that are likely to coordinate lineage specification of the two tissue compartments. This study represents a valuable resource for further experimental examination of foregut organogenesis and the data can be explored via the interactive website https://research.cchmc.org/ZornLab-singlecell.

Prior studies of SM regional identity in the early embryo have been limited. Besides well-known regionalization of Hox gene expression, most studies have largely focused on individual organs such as the gastric or pulmonary mesenchyme ^9, 39, 40^. By comparing single cell transcriptomes across the entire foregut, we revealed an extensive regionalization of the early SM into distinct organ-specific mesenchyme subtypes. It is possible that the divergent transcription signatures of early SM cell types are only transiently utilized to define position and molecular programs during fetal organogenesis. After organ fate is determined, different SM cell types may converge on similar differentiation programs such as smooth muscle or fibroblasts, which are common in all visceral organs. However, our results of fetal SM diversification are interesting in light of the emerging idea of organ-specific stromal cells in adults, such as hepatic versus pancreatic stellate cells and pulmonary specific fibroblasts ^41, 42^. For example, *Tbx4* is expressed in embryonic respiratory SM and later is specifically maintained in adult pulmonary fibroblasts but not in fibroblast of other organs^42^. Future integrated analyses of our data with other scRNA-seq dataset from later fetal and adult organs ^43, 44^ should resolve how transcriptional programs evolve during cellular differentiation, homeostasis and pathogenesis.

One unexpected observation was that the liver bud contained more distinct SM cell states than any other organ primordia with the septum transversum mesenchyme (stm), sinus venosus, two mesothelium and a fibroblast population. This may be due to the fact that unlike other GI organs that form by epithelium evagination, the hepatic endoderm delaminates and invades the adjacent stm, a process that may require more complex epithelial-mesenchymal interactions with the extracellular matrix. Our transcriptome analysis is consistent with lineage tracing experiment showing that the early stm gives rise to the mesothelium, hepatic stellate cells, stromal fibroblasts, and perivascular smooth muscle ^45^. It will be important to determine if other organ buds have a similar elaboration of cell types as they differentiate. Alternatively, mesothelium and fibroblast that originate in the liver may migrate to other organ buds. Indeed mesenchymal cell movement is one confounding limitation of our study and there is good evidence that mesothelium of the liver bud, also known as the proepicardium migrates to surround the heart and lungs ^46, 47^. Going forward, it will be important to use technologies that couple lineage tracing with single cell transcriptomics and live imaging ^48^ to resolve these important questions.

The foregut SM and the cardiac mesoderm are closely related, both arising from the anterior lateral plate mesoderm ^10, 11^. A preliminary cross-comparison of our data with recent single cell RNA-seq studies of the early heart suggests that this common origin is reflected in the transcriptomes ^12^. The developing heart tube is contiguous with the ventral foregut SM (also known as the second heart field), with the arterial pole attached to the pharyngeal SM and the venous pole attached to the lung/liver SM. Fate mapping studies indicate that the second heart field gives rise to heart tissue as well as pharyngeal SM, respiratory SM, and pulmonary vasculature^10, 49^. Indeed, our single cell transcriptomics and genetic analysis of Gli mutants are consistent with previous studies indicating that the epithelium derived HH signals are critical for the development of these cardio-pulmonary progenitors^49, 50^. How the SHF is subsequently segregated into different cardiac and SM lineages is unclear but could be addressed by an integration of our data with other cardiac centric studies.

One important outcome of our study was to use the signaling roadmap inferred from the single cell transcriptomics to direct the development of hPSCs into different SM-like cell types, which to date have been elusive. While more work is needed to optimize the purity of the cell populations and to determine the differentiation potential of each of these cell populations, this system provides a unique opportunity to model human fetal mesenchyme development and to interrogate how combinatorial signaling pathways direct parallel mesenchymal fate choices. These hPSC-derived SM-like tissue may also have important applications for tissue engineering. To date, most hPSC-derived foregut organoids (gastric, esophageal and pulmonary) tend to lack mesenchyme, unlike hindgut derived intestinal organoids. This is because the differentiation protocol needed to make foregut epithelium is not compatible with mesenchymal development. The protocols we have defined here should enable the recombination of DE and SM organoids, an important step towards engineering complex foregut tissue for regenerative medicine. Looking forward, our approach of computational inferring signal interactions coupled with manual curation and a deep understanding of tissue anatomy could be used to predict cell-cell interactions in other organ systems or those that drive pathological states such as the cancer niche.

## Supporting information

upplementary Table S1 Endoderm_Mesoderm_clusters_markers

Supplementary Table S2 signaling metagenes

Supplementary Table S3 HH-Gli regulated transcript

Supplementary Table S4 antibodies_Primers

Supplementary Table S5 RNAscope probes

## ACKNOWLEDGEMENTS

This work was supported by grant NICHD P01HD093363 to AMZ and JWM. TT is a New York Stem Cell Foundation – Robertson Investigator. MM research is supported by a Japanese Grant-in-Aid for Scientific Research (B)(17H04185). KK is supported by a Memorial Foundation postdoctoral fellowship and a RIKEN-CuSTOM collaborative grant for Promotion of Joint International Research (A) (18KK0423). This project was supported in part by NIH P30 DK078392 (Integrative Morphology, Sequencing and Pluripotent Stem Cell and Organoid Cores) of the Digestive Disease Research Core Center in Cincinnati. We are grateful to Minzhe Guo, Yan Xu, and Nathan Salomonis for help with the pseudotime and trajectory analysis, to James Briggs and Caleb Weinreb from the Alon Klein lab for advice on SPRING and members of the Rahul Satija lab for help with Seurat. Drs. Rafi Kopan and Emily Miraldi provided critical discussions and advice.

## AUTHOR CONTRIBUTIONS

AMZ, TT, JMW and HK conceived the project. JMS and HK prepared the mouse foregut single cell samples. PC performed the computational analysis with input from AMZ, LH, HK and KG. AMZ, LH and HK performed manual data curation. LH, TN, KI and PW performed mouse embryo validation. KK, TN, and YL performed RNA-scope analysis. LH, JMW and AMZ devised the differentiation protocol. LH, KK, KI, AE, LH and YL performed the hPSC differentiation experiments. MM and KK shared unpublished data on mouse ESC differentiations. AMZ, LH and PC wrote the manuscript with input from the authors.

## METHODS

### Embryo collection and single cell dissociation

All mouse experiments were performed in accordance with protocols approved by the Cincinnati Children’s Hospital Medical Center Institutional Animal Care and Use Committee (IACUC). No statistical sample size estimates were performed prior to the experiment, sufficient embryos to generate the material need for the experiments were used. No randomization was utilized as no particular treatment was performed in different groups. Timed matings were set up between C57BL/6J mice and the day where a plug was detected was considered embryonic day 0.5. Staging was validated by counting somite numbers E8.5 (5-10 somites; “s”), E9.0 (12-15s) and E9.5 (25-30s) (Fig. 1a, b). We micro-dissected the foregut between the posterior pharynx and the midgut, removing most of the heart and paraxial tissue and excluded the thyroid. At E9.5, we isolated anterior and posterior regions separately, containing lung/esophagus and liver/pancreas primordia, respectively. We pooled dissected foregut tissue from 16, 20, 18 and 15 embryos for E8.5, E9.0 and E9.5 anterior and E9.5 posterior, respectively isolated from 2-3 litters.

Single cell dissociation by cold active protease protocol was performed as described previously^51^. Rapidly dissected C57BL/6J mouse embryo tissues were transferred to ice-cold PBS with 5 mM CaCl_2_, 10 mg/ml of *Bacillus Licheniformis* protease (Sigma) and 125 U/ml DNAse (Qiagen) and incubated on ice with mixing by pipet. After 7 min, single cell dissociation was confirmed with microscope. Cells were then transferred to a 15 ml conical tube, and 3ml ice cold PBS with 10 % FBS (FBS/PBS) was added. Cells were pelleted (1200 G for 5 min), and resuspended in 2 ml PBS/FBS. Cells were washed three times in 5 ml PBS/0.01%BSA (PBS/BSA) and resuspended in a final cell concentration of 100,000 cells/ml for scRNAseq. Single cell suspensions of each stage were loaded onto the Chromium Single Cell Controller instrument (10x Genomics) to generate single-cell gel beads in emulsion. Single cell RNA-Seq libraries for high-throughput sequencing were prepared using the Chromium Single Cell 5’ Library and Gel Bead Kit (10x Genomics). All samples were multiplexed together and sequenced in an Illumina HiSeq 2500. The technician was blinded during the RNA extraction, library preparation and sequencing.

### Immunofluorescence staining, *in situ* hybridization and RNAscope

Mouse embryos were harvested at indicated stages and fixed in 4% paraformaldehyde (PFA) at 4°C for overnight. The fixed samples were washed with PBS 10 min 3 times and the foreguts were micro-dissected when indicated. Embryos or dissected foreguts were then processed as described previously ^52^ by antibody staining (supplementary table.S4) or processed for in situ hybridization.

For RNAscope on mouse tissue, fixed embryos were immersed in 30% sucrose/PBS overnight, embedded in OCT, cryosectioned (12 μm) onto Superfrost™ Plus slides and stored at -80°C overnight. For RNAscope of adherent hPSC culture cells were differentiated on Geltrex-coated u-Slide 8 well (80826, ibid) and fixed in 4% PFA at room temperature for 30min. Cells were dehydrated with ethanol gradient and stored in 100% ethanol at -20°C. RNAscope fluorescent *in situ* hybridization was conducted with RNAscope Multiplex Fluorescent Detection Reagents V2 (323110, advanced cell diagnostics, Inc.) and Opal fluorophore (akoya biosciences) according to manufacturer’s instructions. Detail procedures were listed on Tables S5.

### Pre-processing 10x Genomics raw scRNA-seq data

Raw scRNA-Seq data was processed using CellRanger (v2.0.0 10xGenomics) (https://github.com/10XGenomics/cellranger). Reads were aligned to mouse genome [mm10] to produce counts of genes across barcodes. Barcodes with less than ∼5k UMI counts were not included in downstream analysis. Percentage of reads mapped to transcriptome was ∼70% each sample. The resulting data comprised 9748 cells in E8.5, 9265 cells in E9.0 7208 cells in E9.5 anterior samples and 5085 cells in E9.5 posterior samples.

### Quality control, dimensionality reduction, clustering and marker prediction

Subsequent QC, and clustering was performed using Seurat [v2.3.4] package in R^53, 54^. Basic filtering was carried out where all genes expressed ≥ 3 cells and all cells with at least 100 detected genes were included. QC was based on nGene and percent.mito parameters to remove the multiplets and cells with high mitochondrial gene expression. After filtering 9748, 9265 and 12255 cells were retained in E8.5, E9.0 and E9.5 samples respectively. Global scaling was used to normalize counts across all cells in each sample [scale factor: 10000] and cell cycle effect was removed by regressing out difference between S phase and G2M phase from normalized data using default parameters. We first clustered each developmental stage separately to identify major cell lineages. Approximately 1500 Highly variable genes (HVG) across each population were selected by marking outliers from dispersion vs. avgExp plot. PCA was performed using HVG, and the first 20 Principal Components were used for cells clustering, which then was visualized using t-distributed stochastic neighbor embedding (tSNE). Marker genes defining each cluster were identified using ‘FindAllMarkers’ function (Wilcoxon Rank Sum Test) in Seurat and these were used to annotate clusters based on well-known cell type specific genes.

Cells from all three time points were integrated with Seurat (v3.0) using a diagonalized canaonical correlation analysis (CCA) was used to reduce the dimensionality of the datasets followed by L2-normalization of canonical correlation vectors (CCV). Finally, mutual nearest neighbors (MNN) were obtained which also are referred as integration anchors (cell pairs) to integrate the cells. First 30 CCs (canonical correlation components) were used for clustering and non-linear dimension reduction approaches (UMAP and tSNE) were used to reduce the dimensions and visualize cells in two dimensions.

### *In silico* selection and clustering for definitive endoderm and splanchnic mesenchyme

Definitive Endoderm (DE) clusters (4,448 cells) were defined by the co-expression of Foxa1/2, Cdh1 and/or Epcam, whereas the splanchnic SM (10,097 cells) were defined by co-expression of Foxf1, Vim and/or Pdgfra as well as being negative for cardiac, somatic and paraxial mesoderm specific transcripts. Cells from DE and SM clusters were extracted from each time point and re-clustered using Seurat [v2.3.4] to define lineage subtypes. Prior to re-clustering blood, mitochondrial, ribosomal and strain-dependent noncoding RNA genes were regressed from the data. Dimensionality reduction, clustering and marker prediction steps were performed as described above for each stage. DE and SM cell subtypes were annotated by manual curation comparing the cluster marker genes with over 300 published expression profiles in the MGI database^20^ and our own gene expression validations. DE and SM clusters from all three time points analyzed together using Seurat (v3.0) integration approach explained above respectively.

### Transcription factor code for DE and SM lineages

To identify TFs with enriched expression specific to different DE and SM cell types ‘FindAllMarkers’ function in Seurat [v3.0] was utilized on set of 1623 TFs expressed in the mouse genome [AnimalTFDB] ^55^. Raw counts of TFs were normalized and scaled in Seurat [v3.0]. Cells in clusters served as replicates in finding marker TFs for each lineage. Wilcoxon rank sum test was used in identifying marker TFs. Top 5 marker TFs were then visualized using DimHeatmap function in Seurat(v3.0)

### Pseudo-time analysis of cell populations spatial organization

To examine whether pseudo-time analysis could inform the spatial organization of cells in the continuous sheet of tissue DE or SM tissue a pseudo-time analysis was performed using URD [v1.0] [Farrell, 2018 #354]. Firstly, in order to calculate pseudotime, transition probabilities were calculated for DE and SM cells at each stage using diffusion maps. Then the calcDM function was used to generate diffusion map components and the first 8 components were used to calculate transition probabilities among cells. Next to calculate pseudotime, root cells were fixed to the most anterior clusters based on manual annotation. Starting from root cells a probabilistic breadth-first graph search using transition probabilities was performed until all the cells in the graph have been visited. Multiple simulations were run and pseudotime equaled average iteration that visited each cell in the graph from the root cells. Following functions in URD were used to calculate pseudotime (“floodPseudotime” and floodPseudotimeProcess”). Finally, density distribution of pseudotime was plotted for each cluster/cell-type using plotDists function. Density distribution of pseudotime, ordered clusters similarly to the manually curated order of cell types along the A-P axis.

### SPRING analysis of cell trajectories

To examine cell trajectories across the three time points, we implemented SPRING [v1.0] ^27^, which uses a k-Nearest Neighbors (KNN) graph (5 nearest neighbors) to obtain force-directed layout of cells and their neighbors. To understand transcriptional change across cell states (lineages), first 40 principle components (PC) were learnt from the latest time point E9.5, and this PC space was used to transform the entire data set (E8.5, E9.0 and E9.5). This transformed data was used to generate a distance matrix which then was used to obtain the KNN graph using the default parameters.

### Inferring a cell-state tree by parent-child single cell voting

To visualize the trajectories in a simple transcriptional cell state tree, we used a parent-child single cell voting approach based on the KNN classification algorithm. First, a normalized counts matrix was generated using the distinguishing marker genes from all DE or SM clusters as features at each stage. Marker genes were used as features to train KNN, during which the KNN learns the distance among cells in the training set based on the feature expression. Each cell was classified based on the Seurat cluster assignment. Cells of a later time point vote for their most likely parent cells in the earlier time point as follows: train KNN using E8.5 cells and test by E9.0 cells voting for E8.5 cells. KNN resulted in vote probability for each cell in E9.0 against each cluster in E8.5, which was subsequently averaged for each cluster in E9.0 against each cluster in E8.5. This approach was repeated with E9.5 cells voting for E9.0 parents. The average vote probability for a given cluster was tabulated, normalized for cluster size and represented as a % of total votes in a confusion matrices. The top winning votes linking later time points back to the preceding time point were displayed as a solid line on the tree. Prominent second choices with >60% of winning votes were reported on the tree as dashed lines. We also compared this vote probability with the confusion matrix resulting from the KNN to assess our transcriptional cell-state tree. In more than 99% cases, these two methods resulted in the same first and second choices, thereby validating deduced parent-child relationships.

We validate the cell state tree assertions using pseudotime analysis, Monocle [v3.0.0] ^14^ was deployed on individual lineages/cell states. tSNE was used for Dimensionality Reduction and principle graph was learnt using SimplePPT. All the other parameters were set to default.

### Calculation of metagene profiles

For six of the major paracrine signaling pathways implicated in foregut organogenesis (BMP, FGF, HH, Notch RA and canonical Wnt), we curated a list of all the well-established ligands, receptors and context-independent pathway response genes that were encoded in the mouse genome (supplementary table S2a -excel tab1). We then calculated a “*ligand-metagene”*, “*receptor-metagene*” and “*response-metagene*” profiles by summing the normalized expression of each individual gene for each pathway (e.g.: Wnt-ligand metagene =ΣWnt1+Wnt2+Wnt2b+Wnt3…Wnt10b expression) in each cell and cluster as follows: Assuming that there are x genes in the gene set and n cells. Gene1 has (a1,a2…an) counts, Gene2 has (b1,b2…bn) counts and so on.

<u>Step1:</u> Each gene’s counts were normalized using the gene’s max count across all DE and SM cells (n=14,545 cells) : Gene1_norm = (a1,a2,..an)/max(a1,a2…an)

<u>Step2:</u> Normalized counts of genes were summed up, for each cell, to generate a metagene_v1 with counts across cells: metagene_v1 = Gene1_norm+ Gene2_norm+..+Genex_norm Presuming summed up counts are: m1,m2,..mn

<u>Step3:</u> Summed counts of metagene_v1 were normalized by max counts of the metagene_v1, to create a meta gene profile for each cell : MetaGene = (m1,m2,..mn)/max(m1,m2…mn)

The average Metagene expression profiles for ligands, receptors and response genes in each DE and SM cluster were then calculated in Seurat [v3.0] using ‘AverageExpression’ function. The average expression profile of metagene across all DE and SM clusters were visualized as a Dotplot using Seurat (v3.0). Average Expression of Metagene expression profiles were scaled from -2 to 2 for Dotplot visualization.

### Prediction of receptor-ligand interactions

A given cell type was scored to be expressing enough ligand to send a signal or enough receptor to respond to ligand if the average ligand-metagene or receptor-metagene expression level was ≥ -1 and expressed in ≥ 25% of cells. (Except for the Notch ligand-metagene where expression threshold of ≥ -1.5 was used due to low overall expression in all cells). These thresholds empirically set to be conservative and benchmarked against experimentally validate signaling interactions in DE liver, lung and pancreas from the literature. Furthermore, we determined the likely hood that a given cell population was responding based on the context-independent pathway response-metagene expression level being ≥ -1 and expressed in ≥ 25% of cells. Context-independent response genes are those genes that are known from the literature to be directly transcribed in most cell types that are responding to a ligand-receptor activation.

DE and SM cell clusters of each stage are ordered along the A-P axis consistent with the location of organ primordia *in vivo* with spatially adjacent DE and SM cell types across from one another in the diagram. To assign receptor-ligand interactions for each cell cluster we first determined if a given cluster was responding based on having response-metagene and receptor-metagene levels ≥ -1 threshold. If the responding cluster also expressed the ligand-metagene level ≥ -1, an autocrine signaling was established. For paracrine signaling, we then identified adjacent cell populations, within the same tissue layer and from the adjacent layer that expressed the ligand-metagene above the threshold and then established a receptor-ligand interaction (arrow). The signal strength was calculated as the sum of the ligand-metagene and the response-metagene values. If this value was ≥ 1, the signal was considered “strong”.

### Comparison of bulk RNA-Seq vs scRNA-Seq

Foregut tissue was dissected from E9.5 double mutant *Gli2-/-;Gli3-/-* (n=3) and *Gli2*+*/-;Gli3*+*/-* heterozygous litter mate controls (n=3). Each dissected foregut was separately used for RNA extraction, library preparation and bulk RNA-Seq. These mice were of mixed strains, and the sex of the embryos were unknown. The CSBB [v3.0] (https://github.com/csbbcompbio/CSBB-v3.0) pipeline was used to aligned to mouse genome [mm10] and determine differentially expressed transcripts between the two gene types were obtained using RUVSeq (LogFC ≥ |1| and FDR ≤0.1). Differentially expressed (DE) genes were clustered using hierarchical clustering and visualized in Morpheus (https://software.broadinstitute.org/morpheus/) across samples.

To compare our bulk analysis to scRNA-Seq, we visualized the expression of DE genes across cells in scClusters. We utilized the ‘DoHeatmap’ function in Seurat [v3.0.0]. Cells were arranged according to Anterior/Posterior axis position of their respective clusters and genes were ordered as returned from the clustering order obtained above. We also performed Gene Set Enrichment Analysis (GSEA) [v3.0] ^56^ to examine statistical enrichment of the DE genes in the gut tube SM (respiratory, esophagus, gastric, duodenum), pharynx and liver SM clusters. Normalized counts of genes across cells and up/down-regulated genes from bulk RNA sequencing were used as custom gene sets to perform the GSEA analysis.

### Maintenance of PSCs

Two hPSC lines were used in this study; 1) WA01-H1 human embryonic stem cells purchased from WiCell (NIH approval number NIHhESC-10-0043 and NIHhESC-10-0062) and 2) human iPSC72_3 generated by the CCHMC Pluripotent Stem Cell Facility. Both cell lines have been authenticated as follows: i) Cell identity; via STR profiling by Genetica DNA Laboratory (a LabCorp brand; Burlington, NC), ii) Genetic stability; by standard metaphase spread and G-banded karyotype analysis in CCHMC Cytogenetics Laboratory and iii) Functional pluripotency; cells were subjected to analysis of functional pluripotency by teratoma assay demonstrating ability to differentiate into each of the three germ layer. Both cell lines routinely tested negative for mycoplasma contamination. hPSC lines were maintained on feeder-free conditions in mTeSR1 medium (StemCell technologies, Vancouver, Canada) on six-well Nunclon surface plates (Nunc) coated with Geltrex (ThermoFisher Scientific) and maintained in mTESR1 media (Stem Cell Technologies) at 37°C with 5% CO_2_. Cells were checked daily and differentiated cells were manually removed. Cells were passaged every 4 days using Dispase solution (ThermoFisher Scientific).

### Differentiation of PSCs into mesenchyme

Differentiation of hPSCs into lateral plate mesoderm was induced using previously described methods ^38^ with modifications. In brief, partially confluent hPSCs were dissociated into very fine clumps in Accutase (Invitrogen) and passaged 1:18 onto new Geltrex-coated 24-well plates for immunocytochemistry and 12-well plates for RNA preparation in mTeSR1 with 1uM thiazovivin (Tocris) (Day -1). Next day, a brief wash with DMEM/F12 is followed with Day0 medium 30ng/ml Activin A (Cell Guidance Systems) + 40ng/ml BMP4 (R&D Systems) + 6uM CHIR99021 (Tocris) +20ng/ml FGF2 (Thermo Fisher Scientific) +100nM PIK90 (EMD Millipore) for 24 hours. A basal media composed of Advanced DMEM/F-12, N2, B27, 15 mM HEPES, 2 mM L-glutamine, penicillin-streptomycin is used for this and all subsequent differentiation. On Day 1, a brief wash with with DMEM/F12 is followed with Day 1 medium 1uM A8301 (Tocris) +30ng/ml BMP4+1uM C59 (Cellagen Technology) for 24 hours. For cardiac mesoderm generation, cells are cultured in 1uM A8301+30ng/ml BMP4+1uM C59+20ng/ml FGF2 from Day2 to Day4 (medium changed every day). From Day 4, cells are cultured in 200ug/ml 2-phospho-Ascorbic acid (Sigma) + 1uM XAV939 (Sigma) + 30ng/ml Bmp4 for 3 days. For splanchnic mesoderm generation, cells are cultured in 1uM A8301+30ng/ml BMP4+1uM C59+20ng/ml FGF2 + 2uM RA (Sigma) from Day2 to Day4 (medium changed every day). To further direct regional splanchnic mesoderm, either 2uM RA+40ng/ml BMP4 is used to promote STM fate for 3 days; 2uM RA+ 2uM PMA (Tocris) is used for 2 days, and then 2uM RA+ 2uM PMA + 100ng/ml Noggin (R&D Systems) is used at the last 1 day to promote esophageal/gastric mesenchyme fate; 2uM RA+ 40ng/ml BMP4 + 2uM PMA is used for 2 days, and then 2uM RA+ 40ng/ml BMP4 + 2uM PMA + 1uM CHIR99021 is used at the last 1 day to promote respiratory mesenchyme fate. Medium was changed every day. Similar results were obtained with WA-01 hES cells and human iPSC 72_3.

### Quantitative RT-PCR

Total RNA was prepared from differentiating human ES cells by using Nucleospin kit according to manufacturer’s protocol. Reverse transcription PCR was performed by Superscript VILO cDNA synthesis kit. QuantStudio 5 and 6 were used for qPCR analyses. Primers for qPCR were listed in Supplementary Table 4. Statistics were performed with PRISM8 (GraphPad Software). Significance was determined by one-way ANOVA, followed by Tukey’s test.

### Immunocytochemistry

Cells were fixed with 4% PFA/PBS for 30min at room temperature. After perforation with 0.5% Triton X-100/PBS for 10 min, cells were incubated with 5% normal donkey serum for 2 hours. Cells were incubated with primary antibodies (listed in Supplementary Table 4) overnight at 4°C. Next day, cells were washed with PBS, and then incubated with secondary antibodies for 1 hour at room temperature.

## DATA AND CODE AVAILABILITY

The Source data underlying Figs. 1g,h, 7b,d and Supplementary Figs. S2e-h, S11c,d and S12d are provided as a Source data file. The scRNA-seq and bulk RNA-seq data (including bam, raw counts and cell annotations) are available at Gene Expression Omnibus (GEO): GSE136689 and GSE136687. All the code (scripts, R-packages and software) and their documentation has been uploaded to GitHub [https://github.com/ZornLab/Single-cell-transcriptomics-reveals-a-signaling-roadmap-coordinating-endoderm-and-mesoderm-lineage]. All the deposited code is available to use with GPLv3.0. The scRNA-seq data can be explored at https://research.cchmc.org/ZornLab-singlecell.

## Supplementary Material

**Fig. S1.**
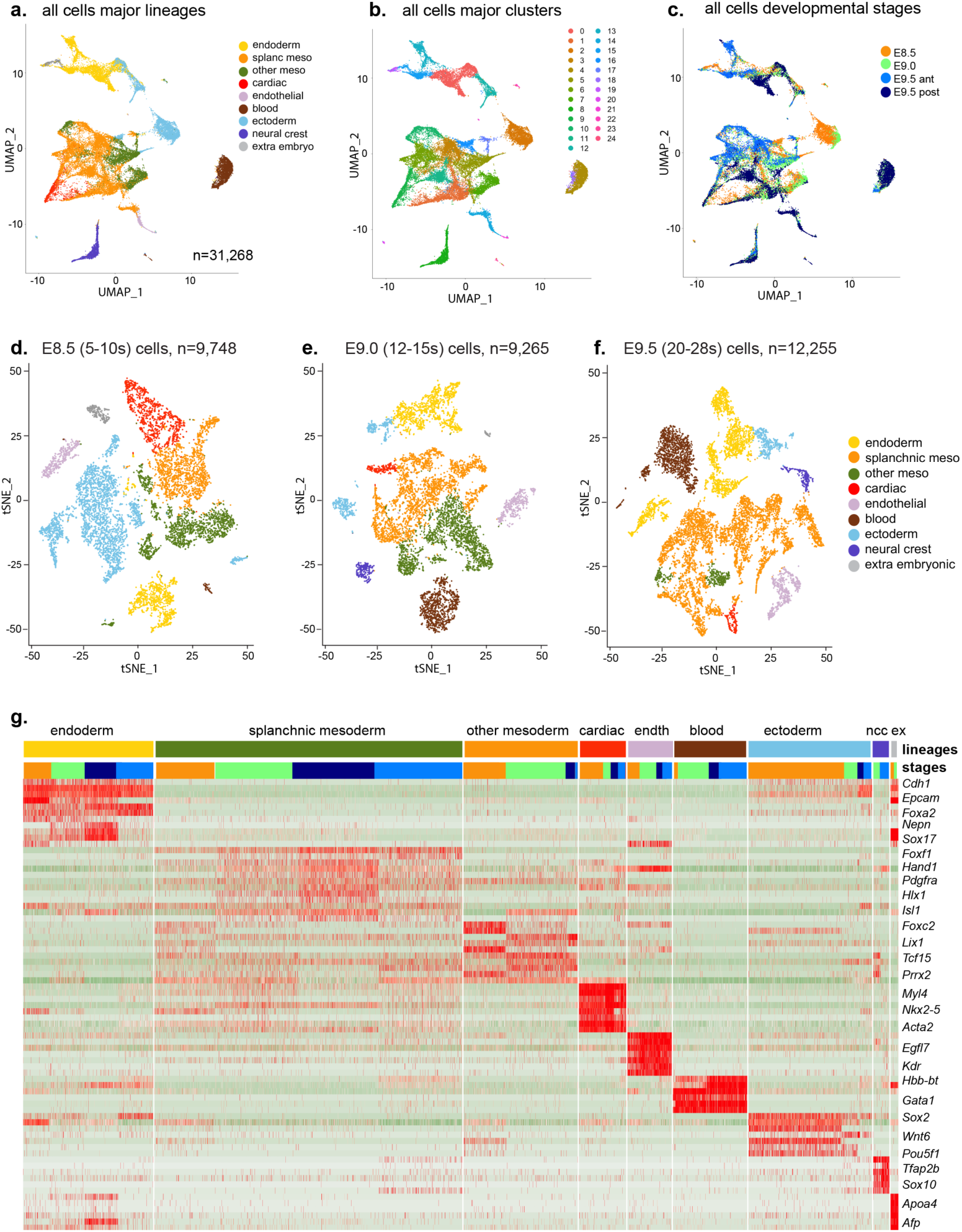
Defining major cell lineages. **a**, UMAP of single cells from all stages with major lineage annotated by known marker genes. **b**, UMAP of all cells from all stages with computationally assigned cell clusters based on transcriptome similarity. **c**, UMAP of all cells from all stages color by stages and regions. **d-f**, tSNE map of single cells from each stage annotated by major lineages at E8.5 in d), E9.0 in e) and E9.5 in f). **g**, Gene expression heatmap of selected markers in individual cells across different lineages and stages.

**Supplementary Fig. S2.**
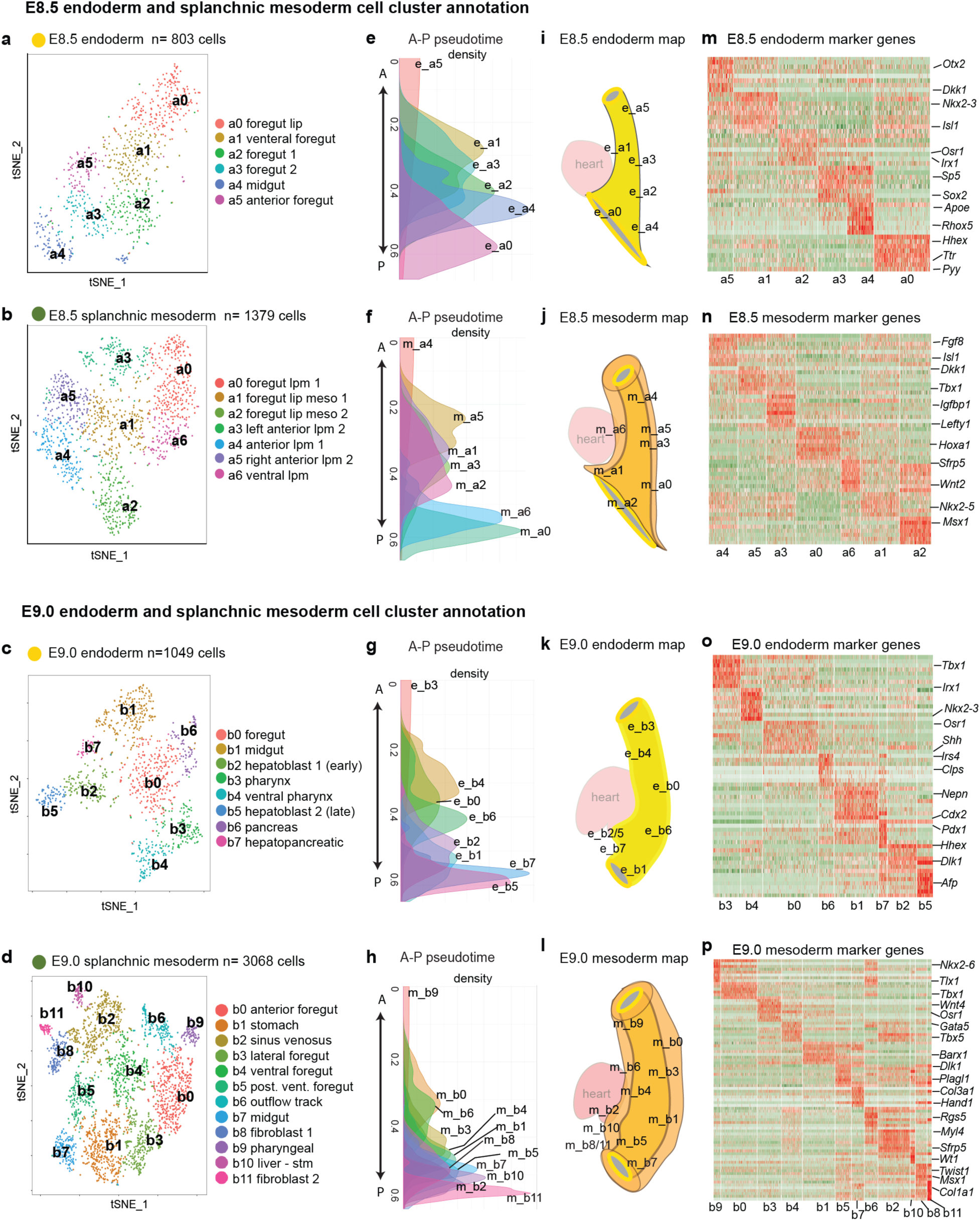
Annotation of E8.5 and E9.0 DE and SM lineages. **a-d**, t-SNE plot of E8.5 DE (**a**), E8.5 SM (**b**), E9.0 DE (**c**) and E9.0 SM cells (**d**) annotations. E8.5 clusters are designated as “a”, E9.0 as “b”, and E9.5 as “c”. **e-h**, Pseudo-spatial ordering of E8.5 DE (**e**), E8.5 SM (**f**), E9.0 DE (**g**) and E9.0 SM cells (**h**) along the anterior-posterior (A-P) axis of the gut tube. **i-l**, Schematics of the mouse embryonic foregut showing the predicted location of E8.5 DE (**i**), E8.5 SM (**j**), E9.0 DE (**k**) and E9.0 SM (**l**) cell types mapped onto the endoderm (yellow) and mesoderm (orange). **m-p**, Heatmap of selected marker gene expression in individual cells across different clusters at E8.5 DE (**m**), E8.5 SM (**n**), E9.0 DE (**0**) and E9.0 SM (**p**).

**Fig. S3.**
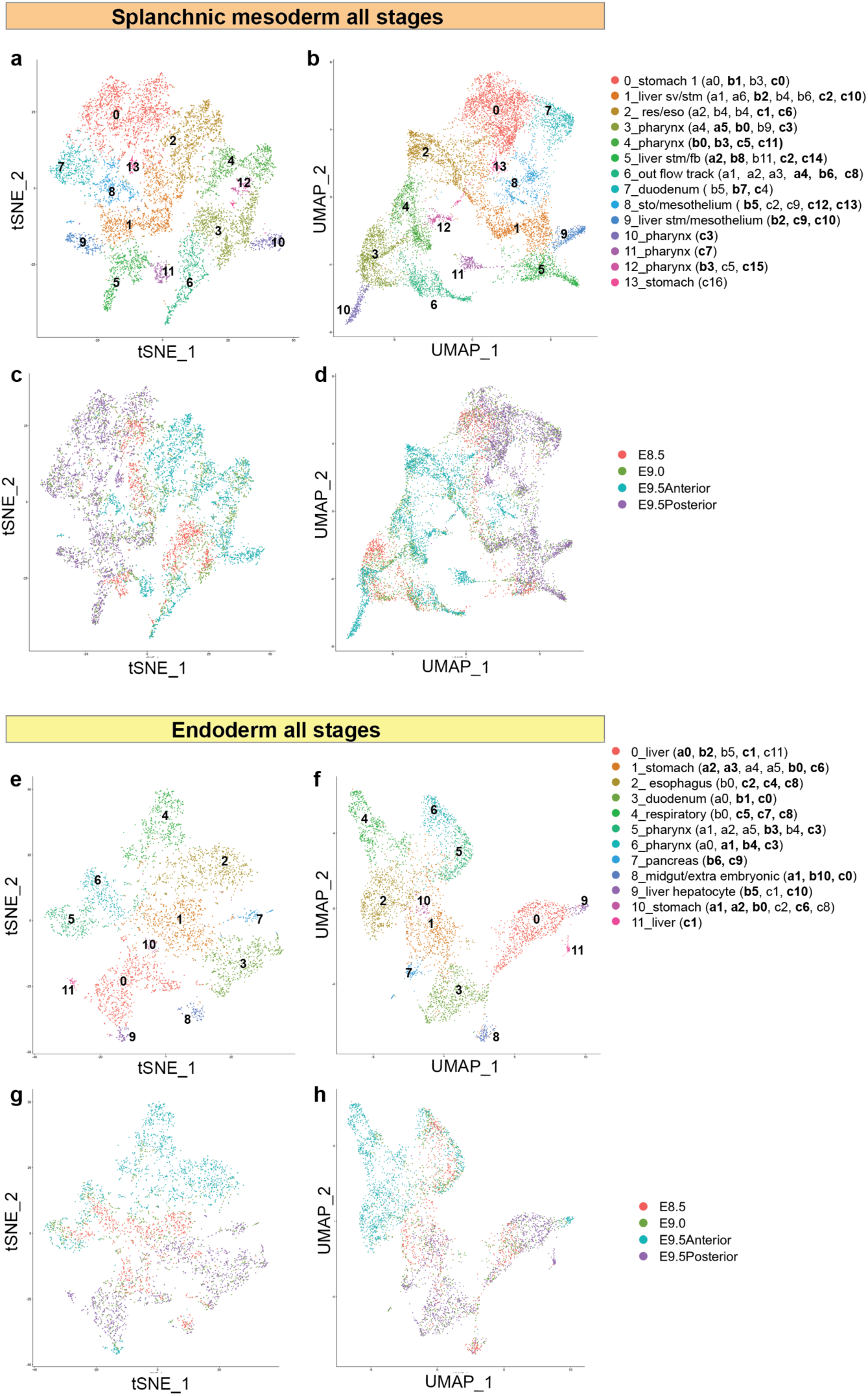
Integrated analysis of DE and SM cells. **a-d**, tSNE and UMAP visualization of all SM cells from all stages annotated by major lineages (a,b) and stages (c,d) **e-h**, tSNE and UMAP visualization of all DE cells from all stages annotated by major lineages (e,f) and stage (g,h). The stage-specific annotations making major contributions to each integrated cluster are indicated in brackets. E8.5 cells = a_clusters, E9.0 cells = b_ clusters and E9.5 = c_clusters.

**Fig. S4.**
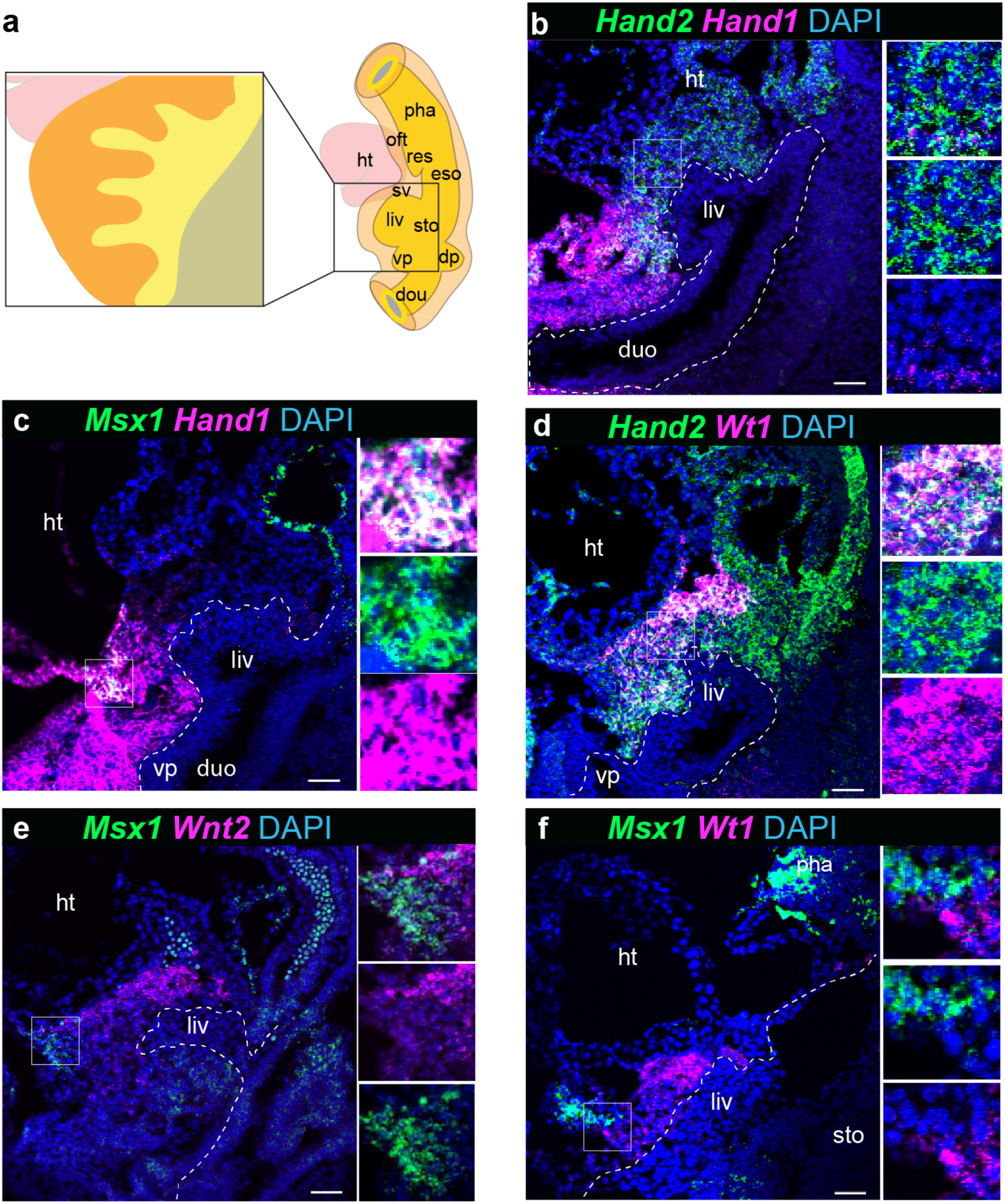
Validation of liver mesenchyme subtypes. **a**, Schematic of mouse embryonic foregut at E9.5. Magnified panel shows sagittal section of the liver bud. **b-f**, RNA-scope *in situ* detection of mesoderm markers on fixed frozen sagittal sections from E9.5 mouse embryos. duo; duodenum, dp; dorsal pancreas, eso; esophagus, ht; heart, liv; liver, oft, outflow tract, pha pharynx, res; respiratory, stm; septum transversum, mesenchyme, sto; stomach, sv; sinus venosus, vp; ventral pancreas. Scale bar 50μm. Insets show merge and separate channels.

**Fig. S5.**
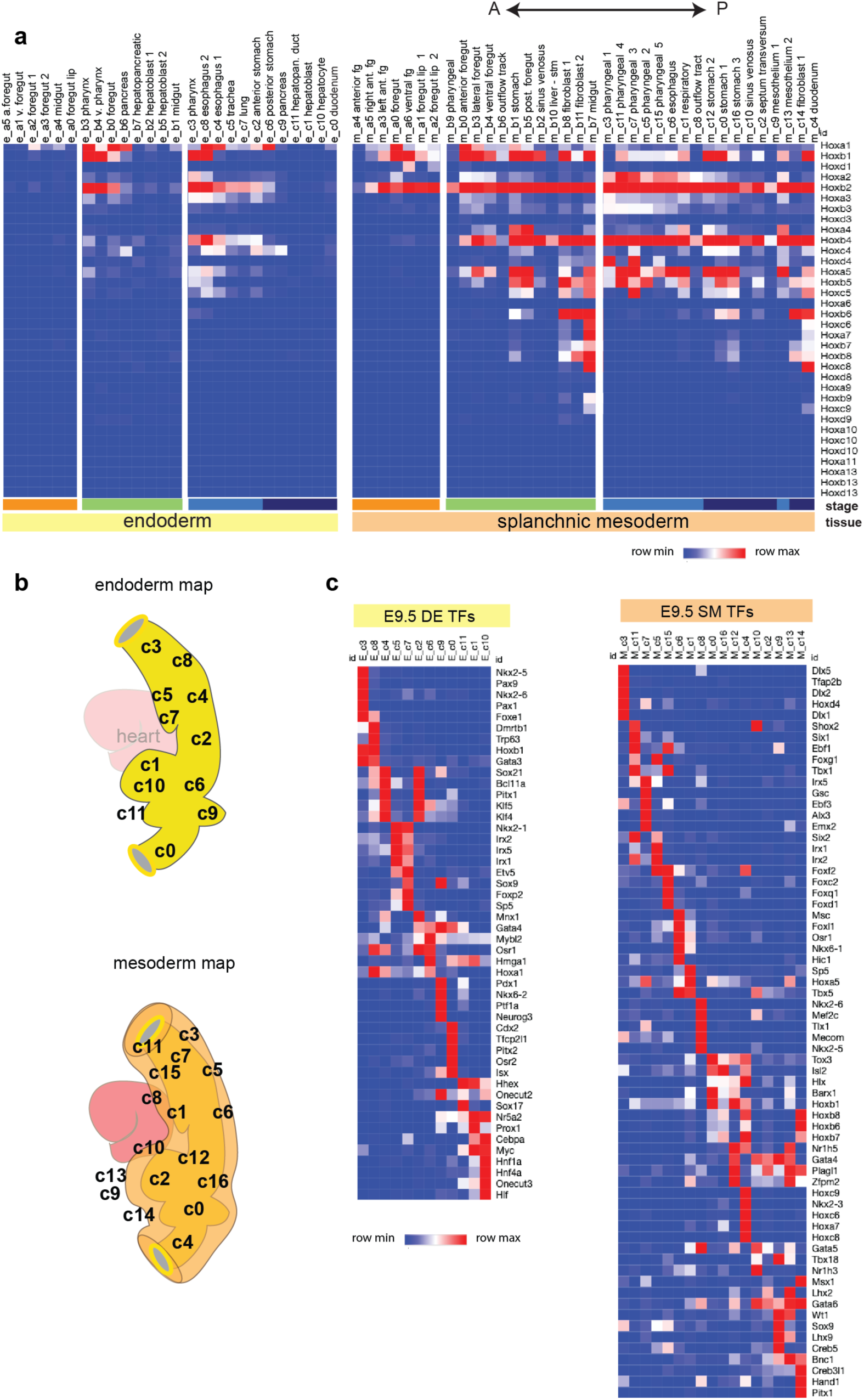
Co-linear *Hox* gene expression and Transcription factor code. **a**, Heatmaps of average *Hox* gene expression across different DE and SM clusters arranged along the A-P axis. Annotations are; E8.5 = a_clusters, E9.0 = b_clusters and E9.5 = c_clusters. **b**, Inferred location of cell clusters in the foregut endoderm and mesoderm. **c**, Transcription factor code. Heatmap showing the average expression of top five distinguishing transcription factor (TFs) differential expressions across E9.5 DE and SM populations. a, anterior; fg, foregut; post, posterior; v, ventral; stm, septum transversum mesenchyme.

**Supplementary Fig. S6.**
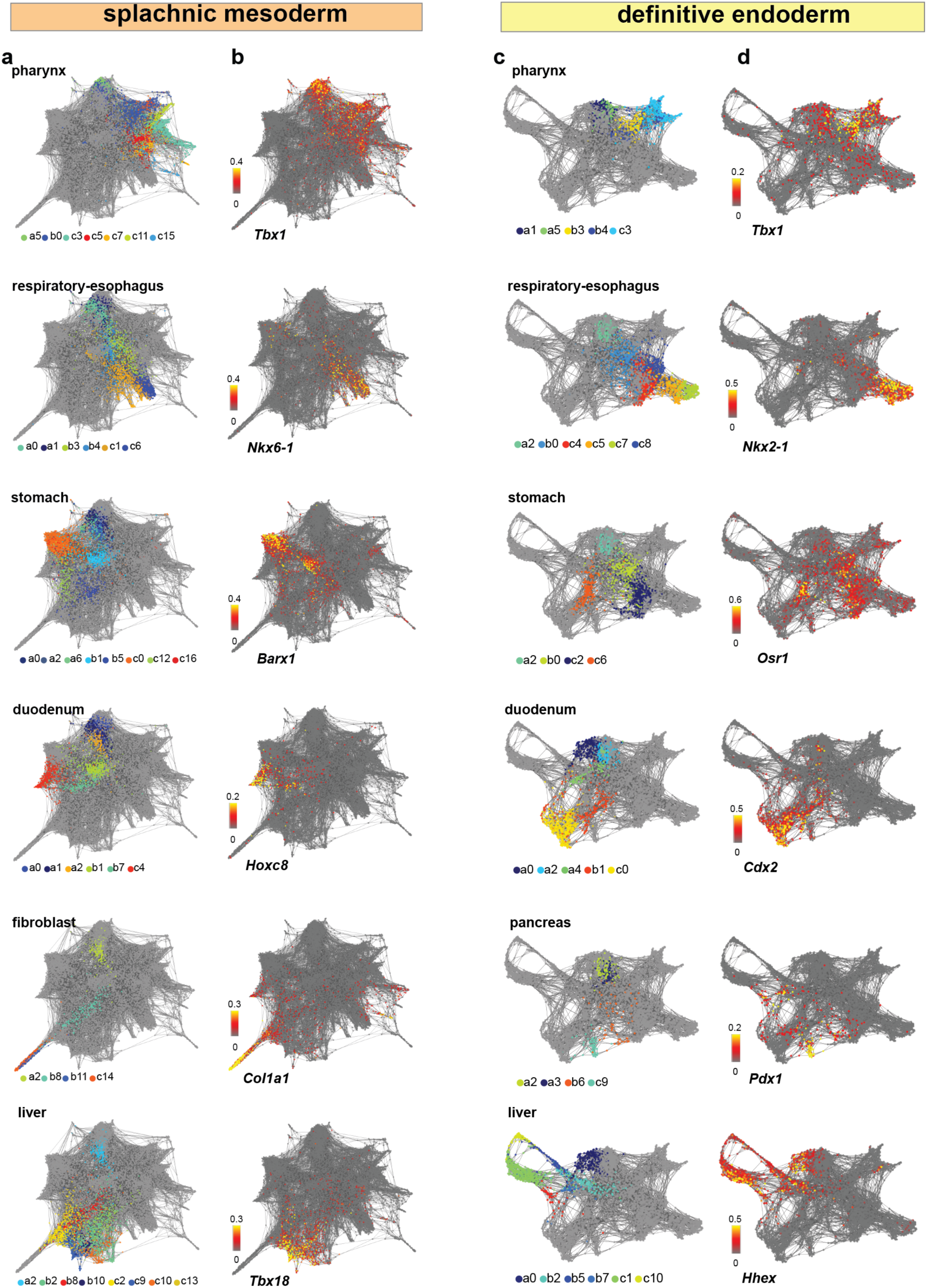
SPRING plot of DE and SM cell trajectories. **a** and **b**, Spring plots of all SM cells (n=10,097) colored by **a**) stage specific lineage annotations and **b**) expression of key marker genes. **c** and **d**, Spring plots of all DE cells (n=4,448) colored by **c**) stage specific lineage annotations and **d**) expression of key marker genes.

**Fig. S7.**
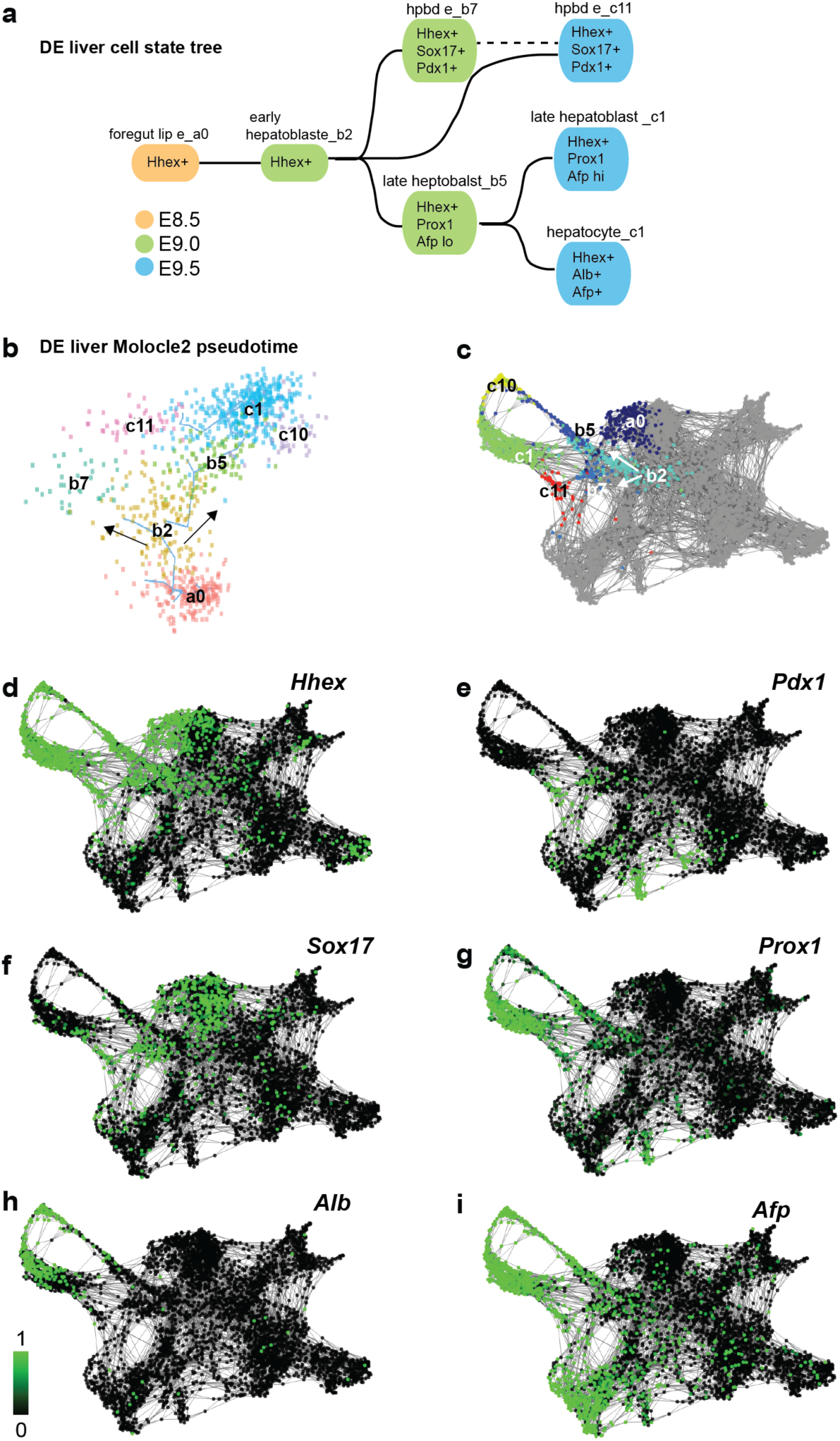
hepatic endoderm development. **a**, Cell state tree of the hepatic endodermal lineage with key marker genes indicated in each cell state. **b**, Pseudotime analysis of the hepatic DE lineage using Molocle_v3 suggests that at E9.0, the e_b2 cluster (early hepatoblasts) is a common progenitor of e_b5 (later hepatoblasts) and e_b7 (hepatopancreatic duct progenitors). **C-i**. SPRING plot with hepatic endodermal clusters colored by **c**) stage specific lineage annotations and **d-i**) expression of key marker genes.

**Supplementary Fig. S8.**
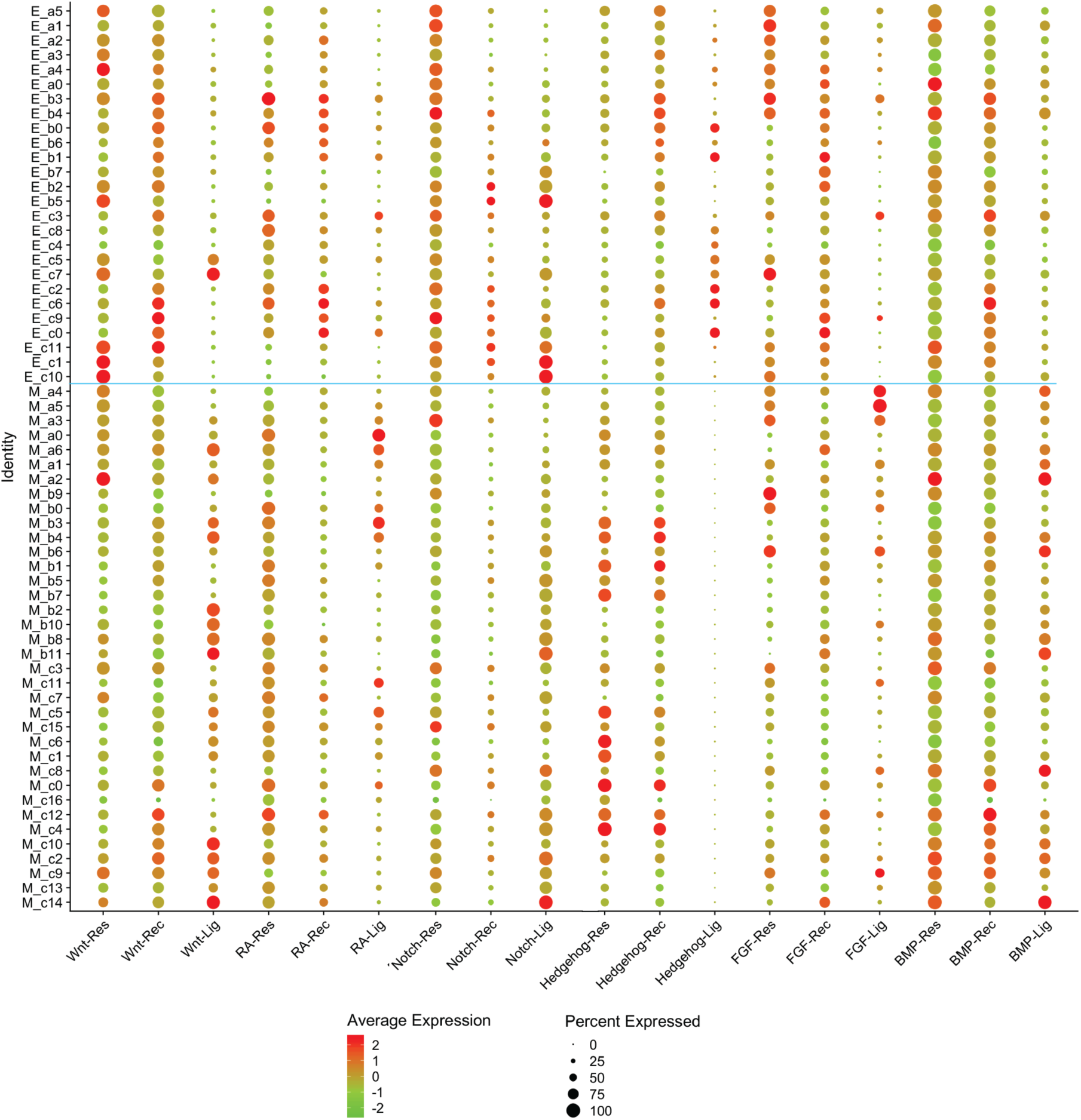
Metagene expression for all ligands, receptors and context-independent response genes. Dot plot showing the average scaled expression (2 to -2) of metagenes (X-axis) in each DE and SM cluster (Y-axis). For each cell signaling pathway (BMP, FGF, HH, Notch RA and canonical Wnt), we calculated a “*ligand-metagene”*, “*receptor-metagene*” and “*response-metagene*” by averaging the normalized expression of each individual gene for each pathway (e.g.: Wnt-ligand metagene =ΣWnt1+Wnt2+Wnt2b+Wnt3…Wnt10b expression/n) in each cell and cluster (see Methods for details). Color and size of the dot represents the metagene expression level in each cluster. (See Supplementary Table S2 for list of genes that make up each metagene and the numeric metagene expression data used to generate the plot).

**Supplementary Fig. S9.**
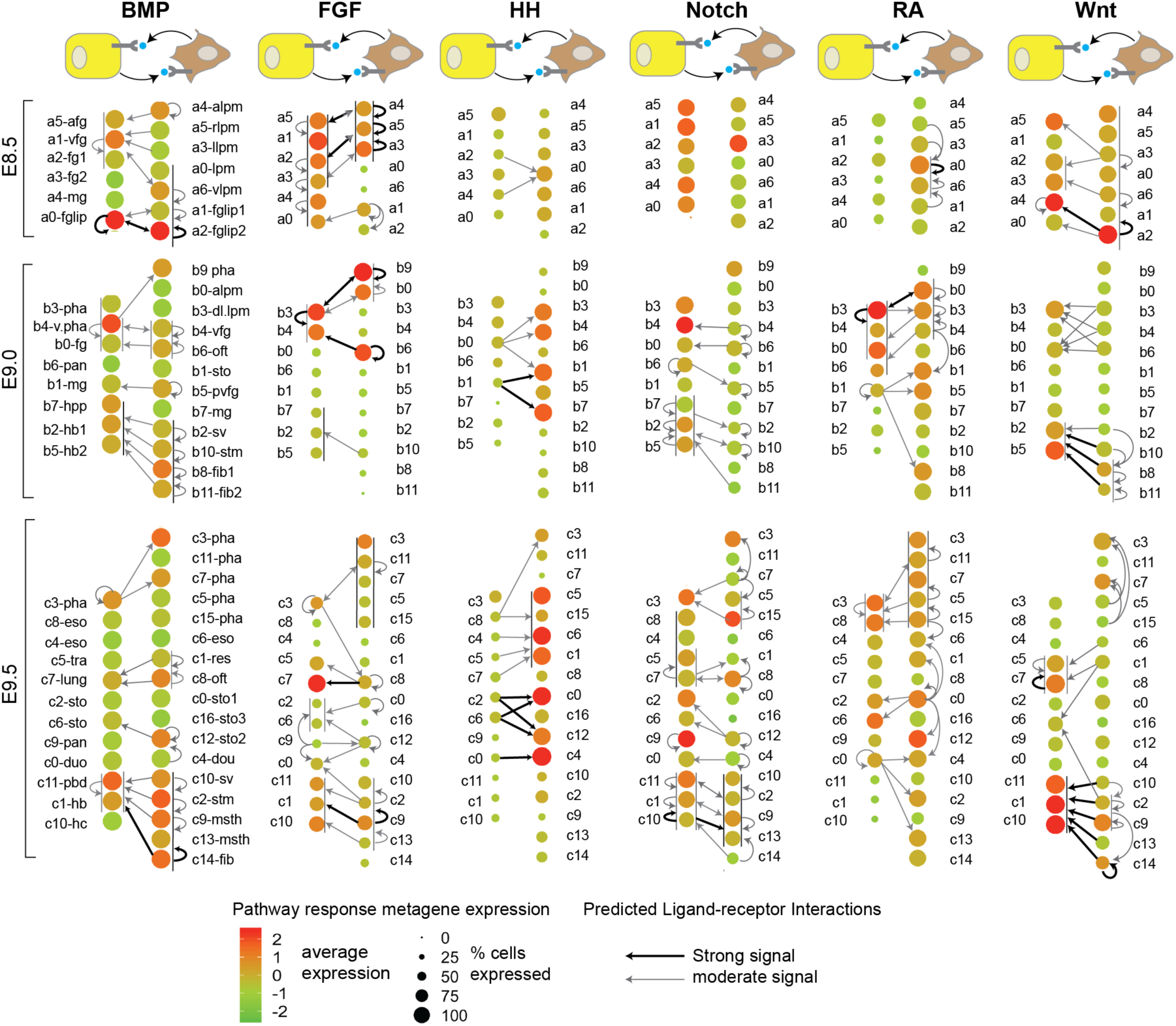
Computationally predicted receptor-ligand interactions between different foregut cell populations. The schematics show paracrine signaling between the DE (yellow cells) and the SM (brown cells) for six major pathways. Below the schematics, DE and SM cell clusters of each stage are ordered along the A-P axis consistent with their location *in vivo*. Spatially adjacent DE and SM cell types are across from one another. Colored circles for each cluster indicate the likelihood that the cell population is responding to the signal based on the pathway response metagene expression levels (see Methods). Arrows represent the predicted source of the ligands showing paracrine and autocrine receptor-ligand pairs inferred from metagene expression profiles. Receptor-ligand pairing (arrows) were restricted to cell populations in close spatial proximity (see Methods for details). Thin vertical lines next to a group of clusters indicate different cell populations in spatial proximity that are all responding similarly.

**Supplementary Fig. S10.**
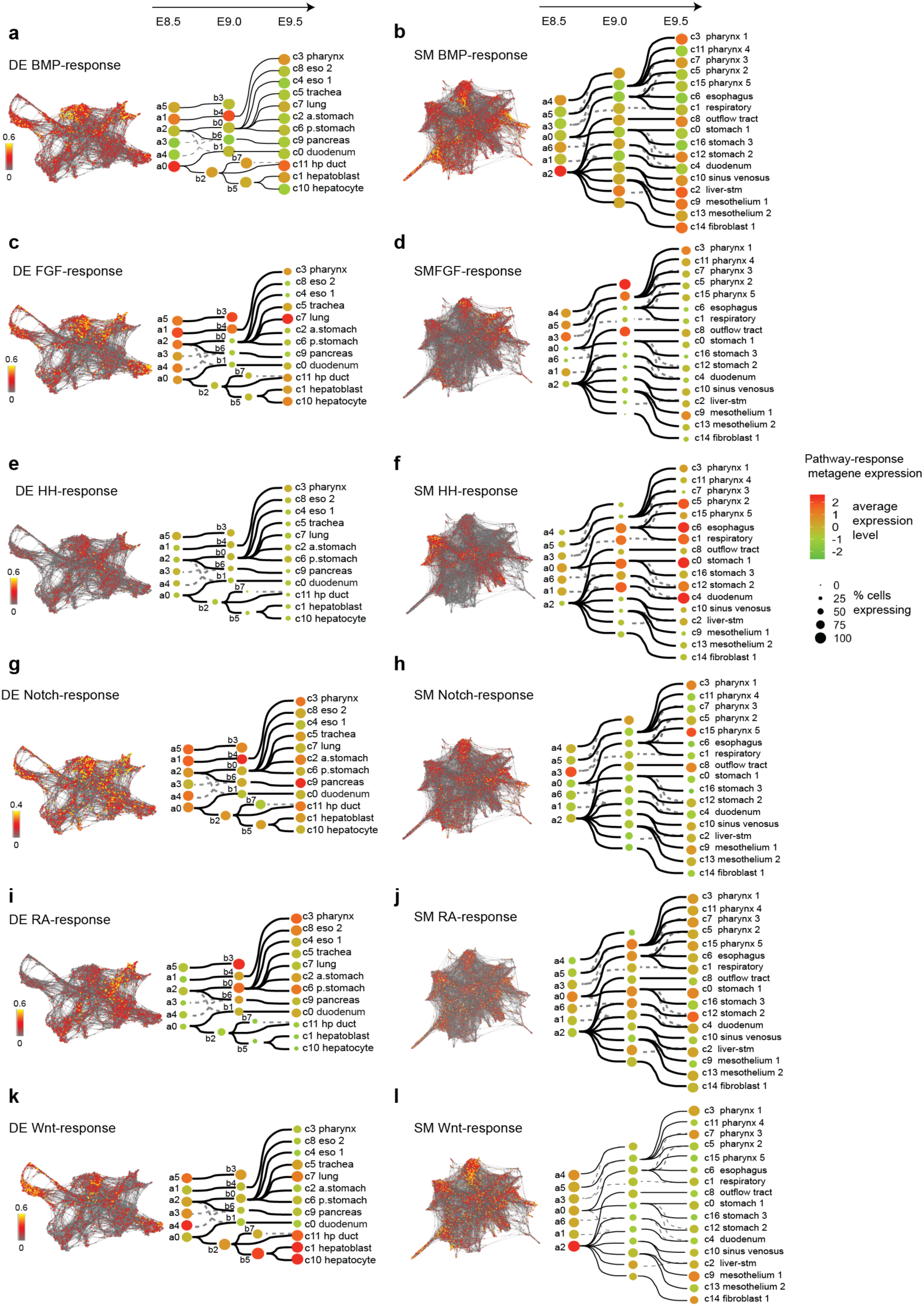
Predicted temporal and spatial dynamics of signaling responses. **a-i**, Expression levels of the pathway response-metagene (From Fig. S9) projected onto the DE and SM SPRING plots and cell state trees for the BMP (a-b), FGF (c-d), HH (e-f), Notch (g-h), RA (i-j) and canonical Wnt (k-l) pathways. This shows how coordinated spatial domains of signaling activity that correspond to cell lineages, are predicted to change over 24 hours from E8.5 – E9.5.

**Supplementary Fig. S11.**
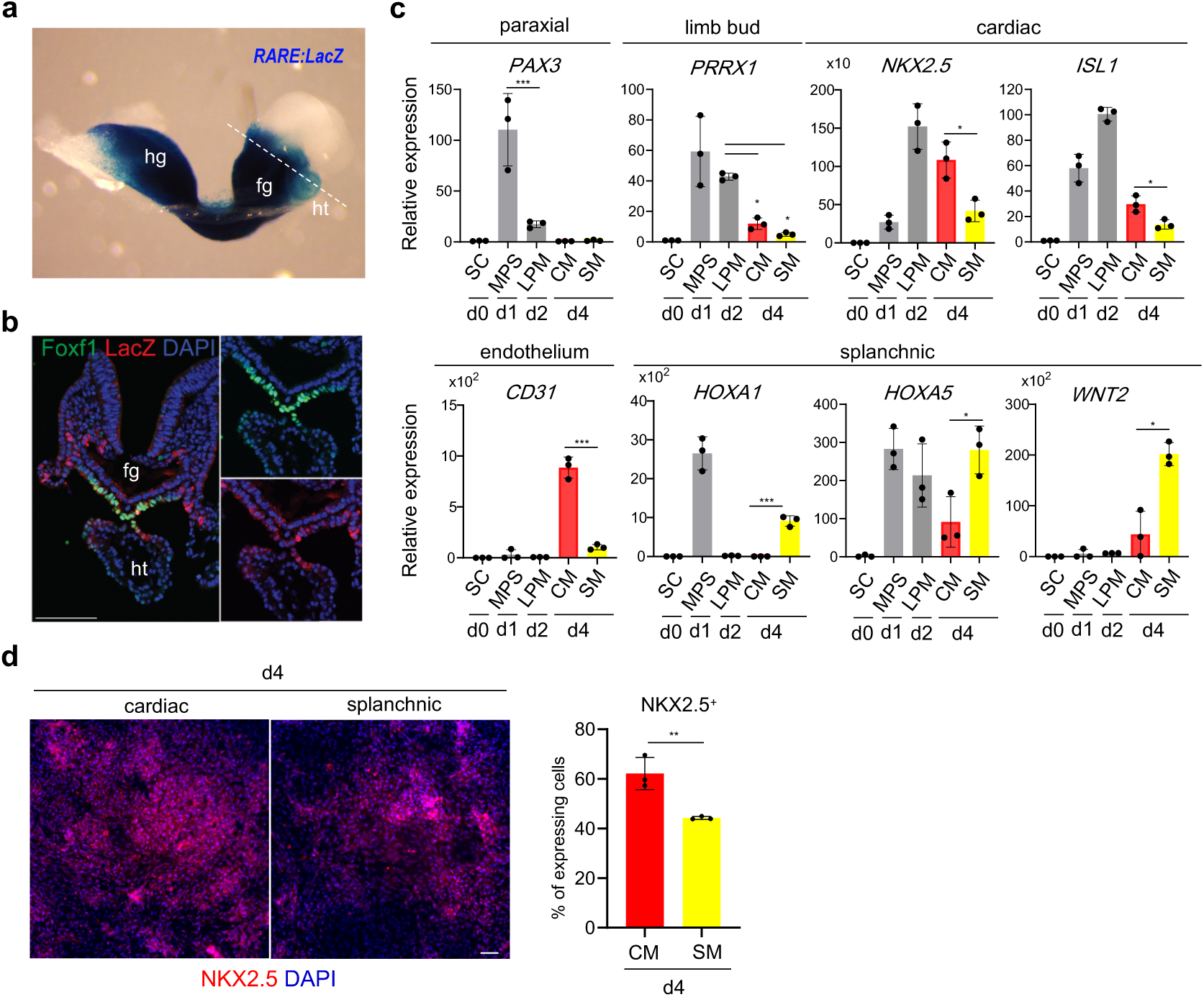
RA suppresses cardiac mesoderm and promotes splanchnic mesoderm progenitors. **a**. Staining of *RARE-lacZ* transgenic mouse embryos confirms the single cell RNA-seq predictions that RA activity is higher in the splanchnic mesenchyme, than the cardiac mesenchyme at E8.5. **b**, Immunostaining of transversal section of *RARE-lacZ* transgenic mouse embryos. **c**. d4 PSC-derived SM cultures assayed by RT-PCR for paraxial mesoderm (*PAX3*), limb bud (*PRRX1*), cardiac mesoderm (*NKX2*.*5, ISL1*), endothelium (*CD31*), and SM (*HOXA1, HOXA5, WNT2*) markers. **c**. Immunostaining of d4 cultures. Scale bar; 50μm. **d**, Quantification of NKX2-5+ cells. fg; foregut, hg; hindgut, ht; heart, SC; Stem Cell, MPS; Middle Primitive Streak, CM, Cardiac Mesoderm, SM; Splanchnic mesoderm, Columns show the means ± S.D. (n=3). Tukey’s test, *p<0.05, **p<0.005, ***p<0.0005. Source data are provided in the Source Data file.

**Fig. S12.**
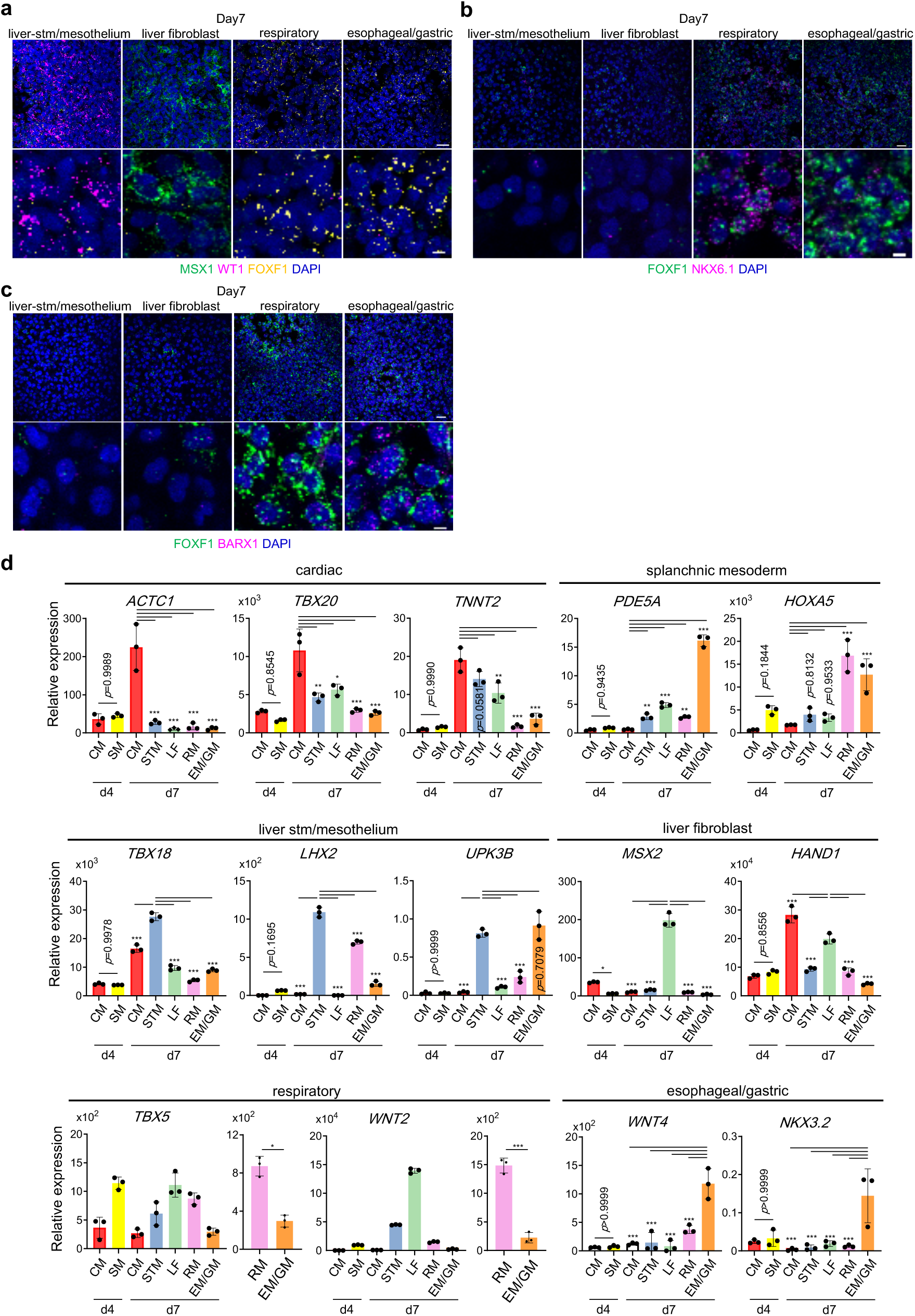
Additional analysis of d7 SM-like PSC cultures. **a-c**, RNA-scope *in situ* analysis of different d7 SM-like cultures. Scale bar 50μm (*Upper panels*), 10μm (*Lower panels*) (quantification in Fig. 7d) **d**, RT-PCR analysis of mesoderm subtype markers based on the mouse scRNA-seq data; cardiac (*ACTC1, TBX20, TNNT2*), early SM (*PDE5A, HOXA5*); liver-stm/mesothelium (*TBX18, LHX2, UPK3B*), liver-fibroblast (*MSX2, HAND1*), esophageal/gastric (*WNT4, NKX3-2*). SC; Stem Cell, MPS; Middle Primitive Streak, CM, Cardiac Mesoderm, SM; Splanchnic Mesoderm, STM; Septum Transversum Mesenchyme, LF; Liver Fibroblast, RM Respiratory Mesenchyme, EM/GM; Esophageal/Gastric Mesenchyme. Columns show the means ± S.D (n=3). Tukey’s test, *p<0.05, **p<0.005, ***p<0.0005. Source data for are provided in the Source Data file.

**Supplementary Table S1. a**, Top distinguishing marker genes for endoderm clusters (excel file tab-1). **b**, Top distinguishing marker genes for splanchnic mesoderm clusters (tab-2). **c**, Transcription factors with enriched expression in endoderm clusters (tab-3). **d**, Transcription factors with enriched expression in splanchnic mesoderm clusters (tab-4).

**Supplementary Table S2. a**. List of BMP, FGF, HH, Notch and canonical Wnt pathway genes used to calculate metagene profiles (tab-1). **b**. Normalized and scaled average expression of each metagene in each DE and SM cluster (tab-2). **c**. Log2 average expression of metagene profiles (tab-3).d. Average Log2 counts of pathway genes in each cluster; BMP (tab-4), FGF (tab- 5), HH (tab-6), Notch (tab-7), RA (tab-8) and Wnt (tab-9).

**Supplementary Table S3**. Differentially expressed transcripts in the bulk RNA-sequencing of E9.5 mouse foregut comparing *Gli2-/-;Gli3-/-* to *Gli2*+*/-;Gli3*+*/-* littermates.

**Supplementary Table S4**. Information on antibodies and RT-PCR primers.

**Supplementary Table S5**. Protocol for RNA-scope *in situ* hybridization of mouse frozen section (tab-1) and human adherent PSC cultures (tab-2).

**Supplementary Movie 1**. Whole-mount staining of mouse embryo foregut at E9.5. Foregut was stained for Nkx2.1 (*red*), Nkx6.1 (*green*), Foxa2 (*blue*), and DAPI (*grey*).

**Supplementary Movie 2**. Whole-mount staining of mouse embryo foregut at E9.5. Foregut was stained for Foxf1 (*red*), Nkx6.1 (*green*), Cdh1 (*blue*), and DAPI (*grey*).

## Notes

https://research.cchmc.org/ZornLab-singlecell

### Competing Interest Statement

The authors have declared no competing interest.

